# Transport properties of canonical PIN-FORMED proteins and the role of the loop domain in auxin transport

**DOI:** 10.1101/2024.02.15.580428

**Authors:** Dorina P. Janacek, Martina Kolb, Lukas Schulz, Julia Mergner, Bernhard Kuster, Matouš Glanc, Jiří Friml, Kirsten ten Tusscher, Claus Schwechheimer, Ulrich Z. Hammes

## Abstract

Indole-3-acetic acid (IAA), the most abundant endogenous auxin is transported in plants in a polar fashion by PIN-FORMED (PIN) transporters and controls virtually all plant growth and developmental processes. Canonical PINs possess a long and largely disordered cytosolic loop domain which is shorter in non-canonical PINs. Auxin transport by canonical PINs is activated loop phosphorylation by kinases. While the structure of the transmembrane domains of these transporters was recently solved, their transport properties remained poorly characterized and particularly the relative roles of the transmembrane and loop domain therein. In this study we used flux studies to obtain quantitative kinetic parameters of IAA transport mediated by canonical PINs as well as of chimeras between transmembrane and loop domains of different PINs upon their activation by D6 PROTEIN KINASE or PINOID. We found that the transporters possess distinct transport properties that are due to both the transmembrane and loop domain. To demonstrate the physiological relevance of these distinct transport properties, we modelled root tip IAA distribution patterns and investigated the potential of different PINs to complement the agravitropic root growth phenotype of the *pin2* mutant when expressed in the *PIN2* domain. We found a strong correlation between transport parameters and physiological output indicating that in addition to PIN polarity a low transport rate in the *PIN2* expression domain is required for gravitropic growth. Overall, the data show that the loop domain is not only required for activation of PIN-mediated auxin transport but has an additional role in the transport cycle by a currently unknown mechanism.

## Introduction

The phytohormone auxin (indole-3-acetic acid, IAA) controls essentially all aspects of plant growth and development. Many developmental effects controlled by auxin require its redistribution within the plant by polar auxin transport ^1–8^. Different types of transporters participate in the transport of IAA into and out of the cell ^9^. Among these, PIN-FORMED (PIN) auxin exporters are the key players in polar auxin transport because several family members are polarly localized in the plasma membranes of many cells, thus providing directionality to IAA export ^9–11^. Mutations in *PINs* cause well-defined phenotypes. Amongst others, the *pin1* mutant displays the eponymous pin-shaped inflorescence ^12^, the *pin2* mutant has agravitropic root growth ^13^ and the *pin3 pin4 pin7 (pin347)* triple mutant is non-phototropic ^14^.

Recently the structures of PIN1, PIN3 and PIN8 were elucidated by cryo-electron microscopy (cryo-EM) ^15–17^. PINs were shown to transport IAA by a crossover elevator uniport mechanism using the membrane potential as a driving force ^18^. PINs form dimers, with each monomer consisting of ten transmembrane helices separated by a hydrophilic loop domain between helices five and six. The PIN monomer can be divided into a scaffold domain, comprising the first two helices of each bundle (M1–M2 and M6–M7) and a transport domain comprising the remaining helices in two three-helix bundles with the characteristic crossover between the broken helices M4 and M9 at the center (M3-M4a/b-M5 and M8-M9a/b-M10) ^18^.

The cytoplasmic loop between M5 and M6 in PINs can be used to non-phylogenetically classify the PINs into long-loop (or canonical) PINs and short-loop (or non-canonical) PINs. In Arabidopsis PINs the long loop ranges between 297 aa in PIN4 and 329 aa in PIN2 while the short loop is 29 aa in PIN5 and 44 aa in PIN8. *Arabidopsis thaliana* PIN6 possesses a loop of intermediate size (250 aa). Unfortunately, none of the recent structures was able to resolve the loop structure except for a short ∼40 aa beta sheet domain directly following M5 in PIN1 and PIN3 ^15,17^. The intermediate PIN6 and short loop PIN8 are able to transport IAA constitutively ^16,19^. In contrast, transport activity of canonical PINs is under the control of AGC1 and AGC3 kinases, suggesting that parts of the long loop are by default autoinhibitory and phosphorylation is required to overcome this inhibition ^16,20,21^. Several AGCVIII kinase mutants display auxin-related phenotypes. Particularly, higher order mutants of *D6 PROTEIN KINASE (D6PK)* and its three closest relatives *(D6PK-LIKE1-D6PKL-3; D6PKs)* as well as mutants of *PINOID (PID)* display undifferentiated inflorescences, similar to *pin1* mutants, suggesting a role in polar auxin transport ^22–24^. At least five serine residues (S1 – S5) within the loop are critical targets for PIN phosphorylation and activation by D6PK and PID ^21^. S1 – S3 are embedded in highly conserved repeated TPRXS sequence motifs whereas the sequence contexts of the S4 and S5 residues are more variable ^25^. Despite the recent progress on the PIN structures, the transport properties of canonical PINs, particularly the impact of the loop on substrate transport and its phosphorylation status remained elusive. In this study we show that canonical PINs display distinct transport properties and that these transport properties differ among PINs. We show that transport properties additionally depend on the identity of the activating kinase.

We further show that the transport properties of the individual PINs, combined with their localization and polarity, critically impact the physiological role of PINs. Finally, our data demonstrate that the loop is not only critical for transport activation, through its interaction with the transmembrane domains it also contributes to the transport process itself and determines transport rates.

## Results and discussion

### Transport properties of canonical PINs

Knowledge about the biochemical parameters of auxin transport for the individual PINs is crucial to be able to improve existing models of auxin transport for more detailed predictions for plant growth and future studies. Several studies have modelled IAA fluxes but in the absence of biochemical parameters assumed identical transport properties for all PINs ^5,26^. Recent data obtained for PIN1, PIN3 and PIN8 found unexpectedly high and divergent *K_d_* values (substrate binding constants) of ∼200 µM, 100 µM and 40 µM, respectively, indicating that PINs possess different affinities for their substrate ^15–17^. Due to the methods applied, these studies addressed substrate binding. How the activating kinases impact PIN transport properties and auxin affinities and transport rates as a function of substrate concentration remained to be determined. We previously established a Xenopus laevis oocyte-based export system that allows determining transport rates precisely by using defined substrate concentrations administered by microinjection ^21,27^. Using this system, we initially determined the IAA transport rates in the linear range for the canonical PINs both without kinase and combined with D6PK or PID in *Xenopus laevis* oocytes at an internal concentration of 1 µM (Suppl. Fig. 1). We found that all canonical PINs were inactive without kinase and were activated significantly (p < 0.01) by PID. In contrast to PIN2, PIN1 and PIN3 were also activated by D6PK (p < 0.01) but the transport rates were significantly different (p < 0.05) between activation by D6PK and PID (Suppl. Fig. 1D-F). Since we found that the transport properties of the phylogenetically closely related and genetically redundant PIN3, PIN4 and PIN7 were more similar to each other than to the other canonical PINs, we limited the subsequent analyses to PIN3 as a representative member (Suppl. Fig. 1C and F-I) ^7,14,28^.

We determined the IAA transport rates mediated by PIN1, PIN2 and PIN3 in response to different internal IAA concentrations between 0.2 – 10 µM, spanning the entire physiological range of IAA ^29^ (Fig. 1 A, D and G). We found that, over this concentration range, all PINs tested were unable to transport IAA in the absence of a kinase and all PINs could be activated by D6PK as well as by PID (Fig. 1A, D and G). Transport rates of PIN3 were four times higher than transport rates of PIN1, which were about twice as high than those of PIN2 (Fig. 1A, D and G). This indicates that canonical PINs possess different transport rates.

**Figure 1.**
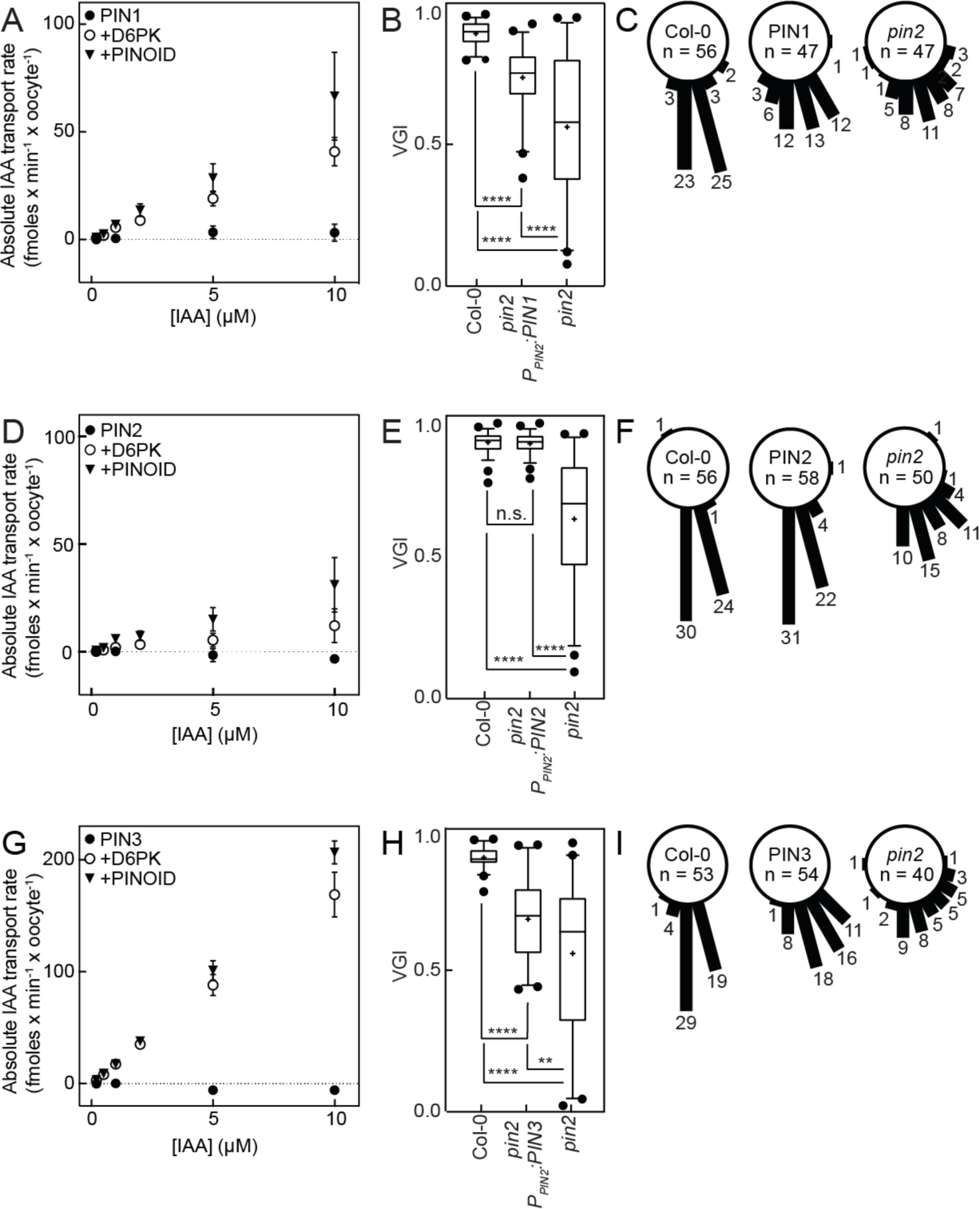
IAA transport properties of canonical PINs in Xenopus oocytes and in plants. **(A)** Transport rates of PIN1 expressed without (●) with D6PK (○) or with PID (▾) as a function of [IAA]_in_. **(B)** VGI of a representative T3 *P_PIN2_:PIN1* line, **(C)** Root angles between root tip and gravity vector of a representative homozygous T3 line of PIN1. **(D)** Transport rates of PIN2 expressed without (●) with D6PK (○) or with PID (▾) as a function of [IAA]_in_. Data are mean and SE of at least n = 3 biological replicates. Data are mean and SE of at least n = 3 biological replicates. **(E)** VGI of a representative T3 *P_PIN2_:PIN2* line, **(F)** Root angles between root tip and gravity vector of a representative homozygous T3 line of PIN2. **(G)** Transport rates of PIN3 expressed without (●) with D6PK (○) or with PID (▾) as a function of [IAA]_in_. Data are mean and SE of at least n = 3 biological replicates. **(H)** VGI of a representative T3 *Pr_PIN2_:PIN3* line, **(I)** Root angles between root tip and gravity vector of a representative homozygous T3 line of PIN3. Data are mean and SE of at least n = 3 biological replicates in A, D and G. Sample sizes are n = 46 in B, n = 56 in D and n = 54 in F comparison to wildtype (n = 55 in B, n = 55 in E, n = 53 in H) and mutant (n = 48 in B, n = 50 in E, n = 40 in H). Box plots range from the 25th to 75th percentiles, whiskers mark the 5th and 95th percentiles and the median is shown, the mean is represented by (+), points below and above the whiskers are individual points. Groups were compared by one-way ANOVA followed by Tukey’s post hoc test. n.s. not significant p-value 0.9861, ** p-value 0.0012, **** p-value <0.0001. Numbers in C, F and I represent individual seedlings.

To rule out that these differences were due to differences in protein levels we determined the amount of transporter, kinase and phosphorylation level in oocytes at the onset of the experiment by liquid chromatography with tandem mass spectrometry (LC-MS-MS) (Suppl. Fig. 2). We found that the abundances of transporters and kinases were very similar and therefore ruled out differences in protein levels as the reason for their different transport rates (Suppl. Fig. 2A-C). We next investigated if the differences are due to PIN phosphorylation. We therefore examined PIN phosphorylation at all serine and threonine residues of the PIN1, PIN2 and PIN3 intracellular loops by phosphoproteomic measurements. Some of the serine residues corresponded to the previously studied S1 – S5, whereas others fit the RXS consensus for putative D6PK and PID target sites ^9^. We found that the kinase-dependent phosphorylation patterns differed between PIN1, PIN2 and PIN3 yet phosphorylation at the critical S1 – S5 residues, with the exception of S2 in PIN1, was in line with these sites being phosphorylated by D6PK and PID ^21,30,31^. Generally, phosphorylation by D6PK and PID was qualitatively and quantitatively very similar on all PINs suggesting that differences in the transport rates between kinases are likely not due to differential phosphorylation (Suppl. Fig. 2 D-F). We found that PIN2 contained the highest number of phosphorylated residues but displayed the lowest transport rates. This indicates that transport rates and number of phosphorylated residues do are not correlated, at least not in our system.

Since we established that both kinases phosphorylate the same residues to a comparable level, it is reasonable to assume that the interaction between the transporter and the kinase impacts the structure of the transporter, thus leading to the observed differences in the kinetics. This demonstrates that canonical PINs do not only display distinct and specific transport properties, but also that these are also subject to modification in a kinase-dependent fashion.

For all PIN/kinase combinations investigated we found a linear correlation between IAA concentration and transport rates (Fig. 1A, D, G). This indicates that PINs possess a very low substrate affinity (high *K_m_*) and that the kinetics at physiological IAA concentrations are below *K_m_* in the seemingly linear range of the Michaelis-Menten curve (Fig. 1A, D and G). For technical reasons such as injection volume, substrate concentration and diffusion from the injection pipette, it is impossible to increase substrate concentration in the oocyte system further but the observation is consistent with the extremely low substrate affinities established independently for PIN1, PIN3 and PIN8 using isothermal titration calorimetry (ITC), surface plasmon resonance (SPR) or solid supported membrane (SSM) electrophysiology, respectively ^15–17^. Since we know from these works that substrate binding can be described by a Michaelis-Menten model, we compared the fit of our data to a linear model versus a Michaelis-Menten model (Extended Data Table 1). We found that, in the case of PIN3 in combination with D6PK, the Michaelis-Menten model was indeed preferred and revealed a *K_m_* of 143.3 µM, albeit with rather low confidence. Nevertheless, this value is in good agreement with the *K_d_* of PIN3 for IAA (160.4 µM) observed by SPR ^15^. For PIN1 and PIN2 with both kinases and PIN3 with PINOID, the linear model was preferred. This indicates that the substrate affinity of PIN3 co-expressed with PID is even lower than that of PIN3 with D6PK. This may also be the case for PIN1 and PIN2, but it is also possible that the comparison was not possible due to the lower transport rates and consequently the lower signal to noise ratio of these transporters compared to PIN3. Together this supports the notion that PINs indeed have substrate affinities that are much lower than the physiological concentration of IAA and operate in the linear transport range.

It remains to be investigated how the interaction between PINs and kinases impacts the transport process in a way that can explain the different transport properties despite similar phosphorylation we observe. A stable heteromer during the transport cycle between transporter and kinase that is an intriguing possibility by which the biochemical properties of a transporter can be modified through physical interaction with a regulating kinase.

### Impact of IAA efflux rates on IAA levels in the root and on root growth

Root gravitropic growth depends on *PIN2* ^13,20,32^. To understand whether the differential transport properties observed in the oocyte system have physiological relevance, we expressed *PIN1*, *PIN2* and *PIN3* under control of a *PIN2* promoter fragment in the *pin2* mutant. We scored the potential of the different PINs to suppress *pin2* agravitropic growth by screening the root angles of 20 independent lines and isolated a representative line (Suppl. Fig. 3A-C). In the homozygous T3 progenies of the representative lines, we compared their vertical growth index (VGI), which considers root length and growth angle of a root to that of the wildtype Col-0, and *pin2* mutants ^33^. We found that, as expected, PIN2 complemented the phenotype to wildtype levels whereas *PIN1* and *PIN3* complemented the phenotype only partially (Fig. 1B, C, E, F, H and I). The difference in VGI between PIN1 and the mutant was higher than between PIN3 and the mutant, indicating that compared to PIN3 PIN1 has more features that are similar to PIN2.

It is well established that PIN2 localizes apically in root epidermis cells and this localization has been suggested to be the reason for proper gravitropic growth ^20,34,35^. To investigate the influence of protein localization we next established that the degree of complementation of the *pin2* mutant was indistinguishable from the wild type proteins and used three independent lines as representative lines of each PIN-GFP fusion construct to determine the polarity index of eight epidermal cells of each line (Fig. 2A and Suppl. Fig. 3D-F). We additionally performed immunolocalizations of the wild type or HA-tagged proteins in epidermal cells (Suppl. Fig. 3G). Consistent with a plethora of previous studies PIN2 displayed the highest degree of apical polarity in epidermal cells whereas PIN1-GFP and PIN3-GFP localized less polarly (Fig. 2A and Suppl. Fig. 3G) ^20,34,36^. The immunolocalizations revealed that PIN1 localized apolarly but predominantly basally whereas PIN3 localized apolarly but with a preference for the apical membrane (Suppl. Fig. 3G). Despite the localization being different from PIN2, PIN1 and PIN3 could partially complement the *pin2* mutant’s VGI indicating that PINs can act redundantly and factors other than localization also must be considered. Interestingly PIN1 complemented the mutant phenotype to a higher degree despite the localization being more apolar which supports the idea that transport rates are critical for complementation. The transport rates we determined for PIN1 and PIN2 were fairly similar and much lower than those determined for PIN3 (Fig. 1 A, D and G). These data indicate that the IAA flux rate through the epidermis, in addition to the proper PIN localization and flux orientation is critical for gravitropic root growth. This is reminiscent of the situation in the protophloem where IAA fluxes must precisely be modulated by the PAX/BRX rheostat module and the efflux rate is critical for cell differentiation ^37,38^. We also investigated the impact of the differential IAA fluxes mediated by the different PIN transgenes on IAA levels in epidermis and cortex cells (Fig. 2B-F). To this end we transformed the representative *pin2, P_PIN2_:PIN* lines described above with the ratiometric auxin sensor R2D2 ^39^. We compared the auxin levels in the cortex and epidermis cell files. Consistent with previous studies, we found that Col-0 seedlings had low auxin in the cortex at both positions and low auxin levels in Q, but high auxin levels at the T position in the epidermis ^40^. In contrast, *pin2* mutants displayed low IAA levels in Q and still low but significantly higher IAA levels in T in the cortex ^40^, while in the epidermis IAA levels in Q were higher leading to no difference between Q and T. As expected, the IAA levels in both cell files of *pin2 P_PIN2_:PIN2* lines matched the pattern observed for wild type plants. For both PIN1 and PIN3, we found that IAA levels were not different between Q and T in either cell file resulting in levels that are intermediate between the *pin2* mutant and the wildtype. This is in line with the observations for the VGI indicating that auxin levels in the cells resulting from export rate are an important factor in addition to proper localization.

**Figure 2.**
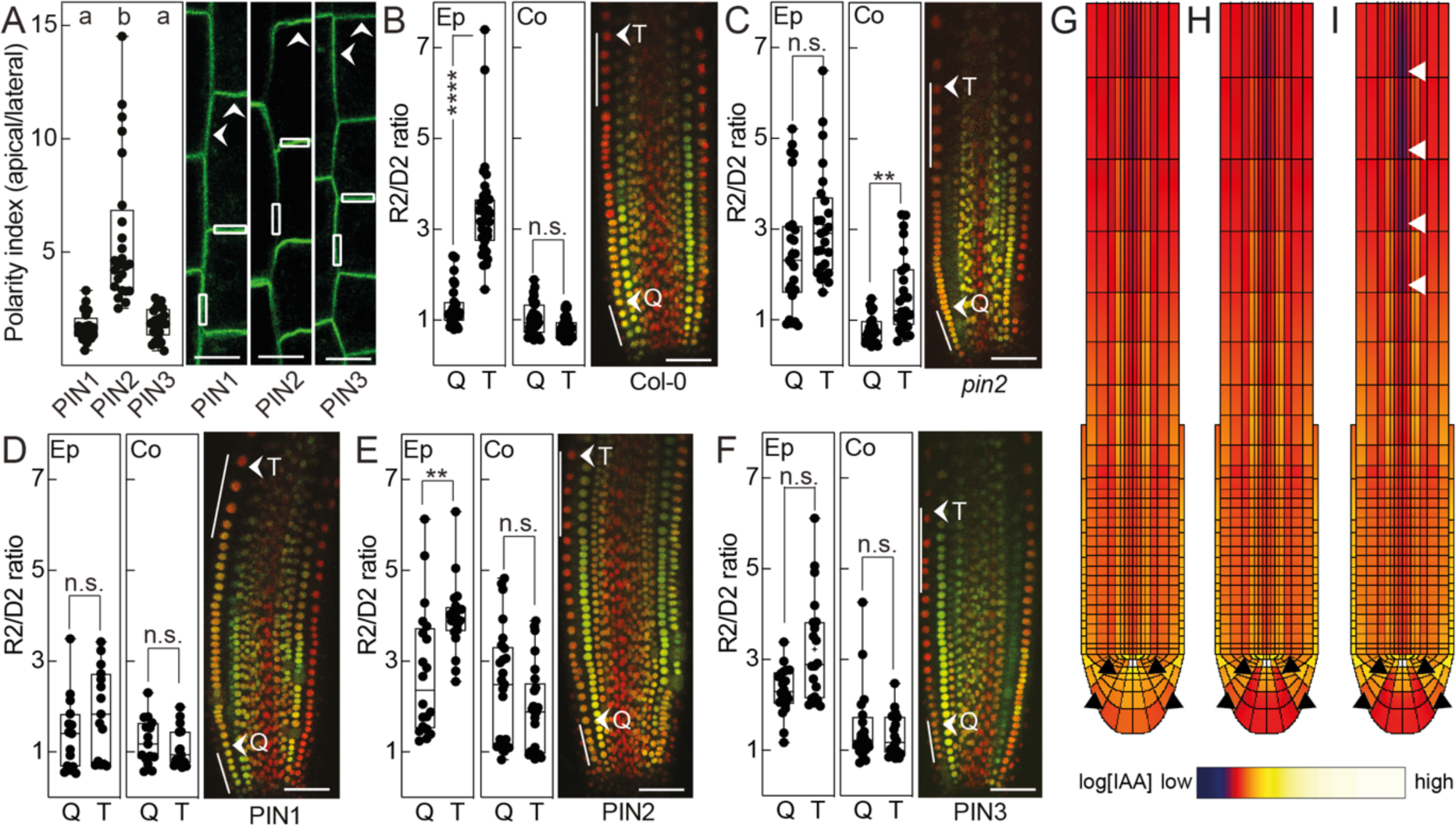
Localization of PINs and IAA response in root tips. **(A)** Polarity indices of *P_PIN2_:PIN1-GFP, P_PIN2_:PIN2-GFP* and *P_PIN2_:PIN3-GFP* in epidermal cells at the onset of elongation in *pin2*. Representative ROIs are indicated by white boxes. Mean intensities of ROIs at the apical and the lateral side of the cells were used to calculate the ratios. Arrows indicate the predominant localization. Scale bars represent 10 µm. Box plots range from the 25th to 75th percentiles, whiskers mark the minimum and maximum values and the median is shown. Groups were compared by a one-way ANOVA, followed by Tukey’s posthoc test (PIN1 vs. PIN2 <0.0001, PIN1 vs. PIN3 0.9592, PIN2 vs. PIN3 <0.0001), n = 24 per genotype. **(B-F)** Ratio of mDII to DII signal in epidermis (Ep) and cortex (Co). **(B)** Col-0. **(C)** *pin2*. **(D)** *P_PIN2_:PIN1-GFP* in *pin2*. **(E)** *P_PIN2_:PIN2-GFP* in *pin2*. **(F)** *P_PIN2_:PIN3-GFP* in *pin2*. The cells indicated by white lines are the first five epidermal cells after anticlinal division of the epidermal/LRC initial cell (Q) and five cells at the transition zone as specified by cell elongation (T) which were measured for individual roots (n = 15-35 for epidermis and n = 15-30 for cortex). Picture shows overlay of DII (green) and mDII (red) signal. Scale bars represent 40 µm. Box plots range from the 25th to 75th percentiles, whiskers mark the minimum and maximum values and the median is shown. Whole data set analyzed by one-way ANOVA, followed by Tukey posthoc test (Col-0: **** p-value <0.0001, n.s. not significant p-value 0.9914. *pin2*: n.s. p-value 0.6432, ** p-value 0.0053. PIN1: n.s. (Ep) p-value 0.9513, n.s. (Co) p-value >0.999. PIN2: ** p-value 0.0075, n.s. p-value 0.0.1938. PIN3: n.s. (EP) p-value 0.1419, n.s. (Co) p-value 0.9986). **(G-I)** Model auxin distribution patterns. The columella is highlighted by black arrowheads**. (G)** Simulated auxin pattern for all PINs with equal transport rates. **(H)** Simulated auxin pattern for 8 times faster transport rates for columella-localized PINs, representing the local presence of faster transporting PIN3. **(I)** Simulated auxin pattern for 8 times faster transport rates in the columella and 2 times faster transport rates in the vasculature (white arrowheads), representing columella localized fastest transporting PIN3 and vasculature localized intermediate fast transporting PIN1.

Taken together, we find that all canonical PINs complement the agravitropic defects of the *pin2* mutant, albeit to different degrees. We further find that PIN1 and PIN2 are very similar in terms of their biochemical transport properties and their potential to complement the *pin2* mutant phenotypes, while differing in polar localization. The phylogenetically more distantly related PIN3 exhibits a higher transport rate and its potential to complement the *pin2* mutant phenotypes is lower than that of PIN1 and PIN2.

### Different transport rates cause different IAA levels in root cells

PIN1, PIN2, and PIN3 act in concert in the Arabidopsis root and form a characteristic auxin maximum around the quiescent center ^10,41^. The concerted activity of these PINs has been modeled in various studies, always under the assumption that all PINs have equal transport rates and all these models predicted a strong auxin accumulation in the columella ^42–44^. This disagrees with the experimentally determined auxin distribution in root cells that found the columella to be relatively free of IAA ^29^. In order to understand whether the different transport rates we determined here lead to a different prediction of the IAA distribution in plants, we used our experimentally determined transport parameters and compared the resulting reparametrized model to the existing model that assumes equal transport rates for all PINs (Fig. 2G-I). We found that using the experimentally determined transport rates the predicted IAA levels in the columella were considerably decreased, and that for this result particularly the higher PIN3 transport rate acting in the columella is essential. This shows that the different transport properties we determined *in vitro* have physiological relevance and can be used to refine models in order to enhance their predictive power.

### Investigating the influence of the loop on PIN transport properties

We were able to show that PINs display distinct transport properties and that these properties are physiologically relevant. It was known from several studies that the loop of the canonical PINs critically determines their localization in the cell ^34,35,45^. It was also suggested that the loop domain and the transmembrane domains of PINs likely coevolved to exert their biological function ^34^. Consistent with the largely disordered structure of the loop the recently published structures of PIN1 and PIN3 could not provide any insight into a possible interaction between the loop and transmembrane regions and the mechanism by which the loop regulates transport activity. To investigate whether the loop domain and the transmembrane domains act in concert and bring about the transport properties of PINs we swapped the loops and the transmembrane domains between PIN1, PIN2 and PIN3 to generate chimeras, e.g., the PIN1-2-1 chimera has the PIN1 transmembrane domains and the PIN2 loop (Suppl. Fig. 4A). We determined the transport rates for the chimeras without kinase or with D6PK or PID at an internal IAA concentration of 1 µM and found that all chimeras mediated IAA efflux from oocytes (Fig. 3A). Consistent with the observations for the wildtype proteins, none of the chimeras mediated IAA transport in the absence of kinase and, like the wildtype proteins, most chimeras displayed higher transport rates upon co-expression with PID than with D6PK. Chimera between PIN1 and PIN2, i.e., PIN1-2-1 and PIN2-1-2, like the wildtype proteins, displayed comparatively low transport rates (Fig. 3A). Transport rates of the chimera between PIN1 and PIN3, i.e., PIN1-3-1 and PIN3-1-1, resembled those of PIN1 more than PIN3, with the transport rates of chimera being in between those of PIN1 and PIN3 (Fig. 3A). Transport rates were further increased in the chimera between PIN2 and PIN3 but still did not reach PIN3 levels. Interestingly, transport rates increased independently of the nature of the transmembrane domain donor. This was very surprising because PIN2 exhibited the lowest transport rates of any PIN but when provided with the PIN3 loop, the chimera showed higher transport rates than the PIN1-3-1 chimera. In summary, the transport data indicate that the loop domain is not merely an on/off switch, but that transmembrane domains and loop domains contribute to the transport properties of the chimeras.

**Figure 3.**
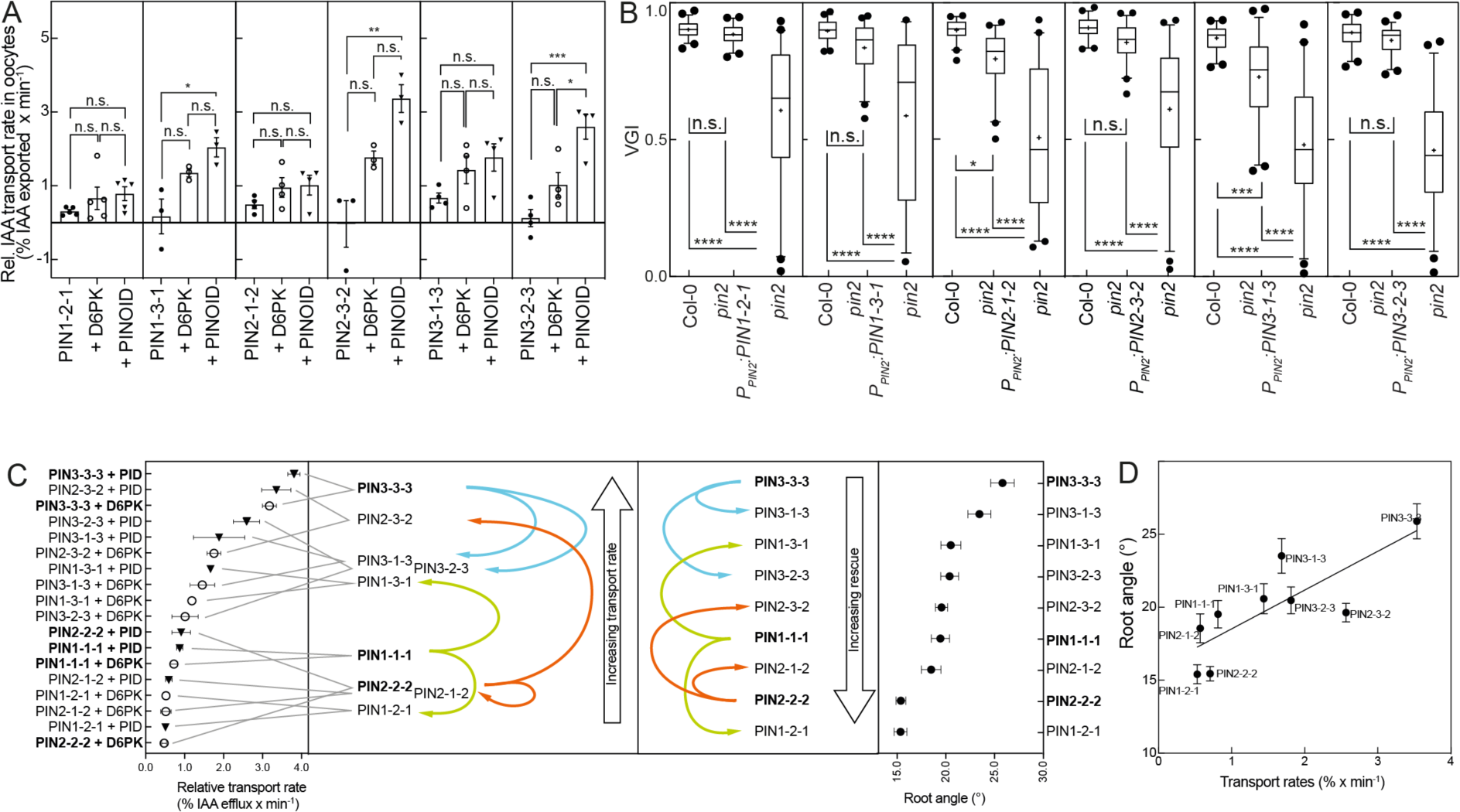
Canonical PIN chimeras are functional IAA transporters whose properties depend on transmembrane and loop donor. (A) Relative IAA export rates for PIN chimeras expressed alone (●) or co-expressed with D6PK (○) or PINOID (▾) in oocytes. Data points are transport rates of n = 3 - 5 biological replicates using oocytes from different females. Groups were compared by one-way ANOVA, followed by Tukey’s posthoc test. PIN2-1-2 vs. PIN2-1-2 + D6PK: n.s. p-value 0.6263, PIN2-1-2 vs. PIN2-1-2 + PINOID: n.s. p-value 0.3433, PIN2-1-2 + D6PK vs. PIN2-1-2 + PINOID: n.s. p-value 0.8402. PIN2-3-2 vs. PIN2-3-2 + D6PK: n.s. p-value 0.0607, PIN2-3-2 vs. PIN2-3-2 + PINOID ** p-value 0.0037, PIN2-3-2 + D6PK vs. PIN2-3-2 + PINOID n.s. p-value 0.0913. PIN3-1-3 vs. PIN3-1-3 + D6PK: n.s. p-value 0.2225, PIN3-1-3 vs. PIN3-1-3 + PINOID: n.s. p-value 0.1001, PIN3-1-3 + D6PK vs. PIN3-1-3 + PINOID: n.s. p-value 0.8104. PIN3-2-3 vs. PIN3-2-3 + D6PK: n.s. p-value 0.1454, PIN3-2-3 vs. PIN3-2-3 + PINOID: *** p-value 0.0007, PIN3-2-3 + D6PK vs. PIN3-2-3 + PINOID: * p-value 0.0129. **(B)** VGIs of the homozygous T3 lines (n = 49 for *P_PIN2_:PIN1-2-1*, n = 45 for *P_PIN2_:PIN1-3-1*, n = 41 for *P_PIN2_:PIN2-1-2*, n = 58 for *P_PIN2_:PIN2-3-2*, n = 46 for *P_PIN2_:PIN3-1-3* and n = 51 *P_PIN2_:PIN3-2-3*) compared to wildtype (n = 49, n = 42, n = 40, n = 57, n = 46, n = 40) and mutant (n = 40, n = 30, n = 40, n = 57, n = 46, n = 56). Box plots range from 25th to 75th percentiles, whiskers mark the 5th and 95th percentiles and the median is indicated. Points below and above the whiskers are drawn as individual points. The mean is indicated by (+). Groups were compared by one-way ANOVA followed by Tukey’s posthoc test (PIN1-2-1: n.s. not significant p-value 0.7753, **** p-value <0.0001. PIN1-3-1: n.s. 0.1919, **** p-value <0.0001. PIN2-1-2: * p-value 0.0140, **** p-value <0.0001. PIN2-3-2: n.s. 0.1123, **** p-value <0.0001. PIN3-1-3: *** p-value 0.0002, **** p-value <0.0001. PIN3-2-3: n.s. 0.6049, **** p-value <0.0001). **(C)** Chimeras ranked by transport rate (left side) and root angles (right side) of the segregating T2 lines. PIN1 transmembrane domain chimeras are represented by green arrows, PIN2 transmembrane domain chimeras are represented by orange arrows and PIN3 transmembrane domain chimeras are represented by blue arrows **(D)** Correlation (r = 0.8) between transport rates in oocytes and root angle of the complementing lines.

In order to test if the transport data derived in oocytes are reflected *in planta* and to be able to compare the chimeras to the wildtype parent domain donors, we next investigated the potential of the chimeras to complement the aberrant VGI of the *pin2* mutant (Fig. 3B and Suppl. Fig. 4B-G). To investigate the localization of the chimera *in planta* we generated GFP-tagged versions of the chimera (Suppl. Fig. 5G-K). We successfully recovered a GFP-tagged line for all chimeras except PIN2-1-2 despite several attempts. All other GFP-tagged chimeras were similar to the untagged versions with respect to the complementation of the mutant phenotypes (Suppl. Fig. 5G-K). We found that the PIN1-2-1 chimera complemented the mutant phenotype to wildtype levels (Fig. 3B and Suppl. Fig. 4B). The chimera also localized apically (Suppl. Fig. 4B) and auxin levels in epidermis and cortex were like wildtype (Suppl. Fig. 4B and Fig. 2B). The PIN1-3-1 chimera also complemented the mutant phenotype to almost wildtype levels (Fig. 3B and Suppl. Fig. 4C) - albeit the protein localized apolarly (Suppl. Fig. 4B). The PIN2-1-2 chimera complemented the mutant partially, but the VGI was significantly different from wildtype (Fig. 3B). Nevertheless, IAA levels in the epidermis were like wildtype (Suppl. Fig. 4D). The PIN2-3-2 chimera localized apically (Suppl. Fig. 4E) and complemented the mutant phenotype to wildtype levels (Fig. 3B). This, however, was the only case were localization and complementation were not reflected by the IAA levels in the epidermis cell file reverting to the wildtype situation (Suppl. Fig. 4E). The PIN3-1-3 chimera localized apolarly (Suppl. Fig. 4F) and complemented the mutant partially (Fig. 3B). Finally, the PIN3-2-3 chimera localized polarly (Suppl. Fig. 4G), complemented the mutant phenotype to wildtype levels (Fig. 3B) and showed the wildtype situation of auxin distribution in the epidermis cell file (Suppl. Fig. 4G). Taken together this indicates that polar localization is clearly correlated with the PIN2 loop as indicated by the localization of the PIN1-2-1 and PIN3-2-3 chimera (Suppl. Fig. 4B and G). This is consistent with previous studies that showed that the Arabidopsis PIN2 loop is sufficient to provide positional cues for Arabidopsis PINs and for other family members in vascular plants ^34,35^. However, additional features must be associated with the PIN2 transmembrane domain as also the PIN2-3-2 mutant localized apically. Nevertheless, the degree of complementation is, like in the case of wildtype proteins, not strictly correlated with polarity as other chimeras, e.g., PIN1-3-1, also complemented the mutant phenotype to wildtype levels.

This supports the notion that lower transport rates from epidermis cells and the auxin ratio, higher in T than in Q position, in the epidermis cell file are more important for gravitropic growth than PIN localization. In order to correlate transport rates and degree of complementation, we ranked the transport rates we measured in the oocytes system and the root angle we observed in the representative complementing line (Fig. 3C). This clearly illustrates that the PIN3 loop increases transport rates of the PIN1 and PIN2 transmembrane domain context above the transport rates of the wildtype proteins. *Vice versa* introduction of the PIN1 or PIN2 loop decreases the transport rates of the PIN3 transmembrane context below the transport rates of the wildtype protein. Exchanging the loops between PIN1 and PIN2 had only minor effects. This correlates well (r = 0.8) with the effect the chimeras have on the root angle *in planta* (Fig. 3D). The lines that contain the PIN3 loop show higher root angles, i.e., a decreased potential to rescue the mutant phenotype in the PIN1 and PIN2 transmembrane domain context. *Vice versa* introduction of the PIN1 or PIN2 loop decreases the root angle i.e., it increases the potential to rescue the mutant phenotype in the PIN3 transmembrane domain context. Like in the transport assays exchanging the loops between PIN1 and PIN2 had only small effects. These data show that the root angle critically depends on the transport rate and, like in the case of the wildtype proteins, lower transport rates in the PIN2 expression domain lead to more gravitropic root growth than higher transport rates.

### Effect of the transport rates on the response to a gravitropic stimulus

So far, we investigated how the different transport rates of the individual PINs affect gravitropic root growth under steady state conditions, showing that different PIN transport properties lead to differential root angles under these conditions. In order to test if the different transport rates also lead to differences in the response to a gravitropic stimulus we modelled the expected IAA distribution under this condition (Fig. 4A). For the *pin2* mutant the model predicted strong IAA accumulation in the root meristem, with a limited auxin asymmetry in the elongation zone, particularly evident for the vasculature, consistent with this being insufficient to drive root bending ^20^. In contrast, for the wildtype, the model predicted a significant IAA asymmetry in the elongation zone of the root (Fig 4A, arrow heads), with clear elevation in epidermal and vascular auxin levels at the lower side of the root consistent with experimental observations and leading to well established root bending ^20,46,47^. The model predicted lower IAA levels as well as a decreased elongation zone IAA asymmetry between the upper and lower side of gravistimulated roots for *PIN1* expressed in the *PIN2* domain and even further reduced IAA levels and asymmetry for *PIN3* expressed in the *PIN2* domain as compared to the wildtype situation (Fig. 4A, arrow heads). Based on these auxin patterns a decreased response to the gravitropic stimulus is expected. To test this prediction experimentally, we turned the plates by 90° and monitored the response in the dark for 16h plates with sucrose (Fig. 4B and C). In line with the prediction, we could confirm that roots expressing *PIN1* in the *PIN2* domain showed the expected delayed response to the stimulus, but the root had completely turned after 16h. Roots expressing *PIN3* in the *PIN2* domain showed an even more delayed response and had not fully adjusted to the gravity vector after 16h. Consistent with previously published data, PIN2 complemented the mutant to wildtype levels ^20,48^. In contrast and as expected *pin2* roots failed to reorient to the gravitropic vector ^20,48^. In order to test the transcriptional response to IAA in the root tips we introgressed the *P_DR5_:GUS* construct into the representative *pin2, P_PIN2_:PIN* lines. Seven hours after the gravitropic stimulus, corresponding to the time point at which we observed the largest differences between the lines, wildtype and *P_PIN2_:PIN2* plants showed low DR5 response in the quiescent center and columella, whereas *P_PIN2_:PIN1* and *P_PIN2_:PIN3* plants displayed high DR5 response in these tissues, as did the *pin2* mutant (Fig. 4C). This indicates that also in this regard PIN1 and PIN3 are similar to each other but different from the wildtype as well as the mutant conditions. Finally, we tested the response of the chimera to the gravitropic stimulus. In order to see more subtle differences we performed these experiments on plates containing sucrose to sustain root growth in the dark as had been suggested recently (Suppl. Fig. 6)^49,50^. In all cases we observed that the *pin2* mutant displayed negative gravitropic growth following the gravitropic stimulus consistent with previous observations under these conditions (Suppl. Fig. 6A-D)^51^. For the wildtype PIN versions, the results of the conditions without sucrose described above are reproducible (Suppl. Fig. 6A). Consistent with the steady state growth conditions, we found that all chimera containing PIN2 transmembrane domain or loop domain also complemented this aspect of the *pin2* mutant phenotype fully. This is consistent with our conclusion that, in addition to proper localization, a low transport rate in the epidermal cell file is required for gravitropic root growth and that the PIN2 transmembrane domain can function essentially like wildtype PIN2 when provided with a loop from PIN1 or PIN3. The findings are also consistent with the idea that the PIN2 gradient observed after a gravitriopic stimulus dampens the IAA gradient and keeps it from becoming too steep ^47^. This is reminiscent of the rheostat model according to which a precisely controlled flux rate rather than absolute levels of IAA control proper protophloem development ^37,38^.

**Figure 4.**
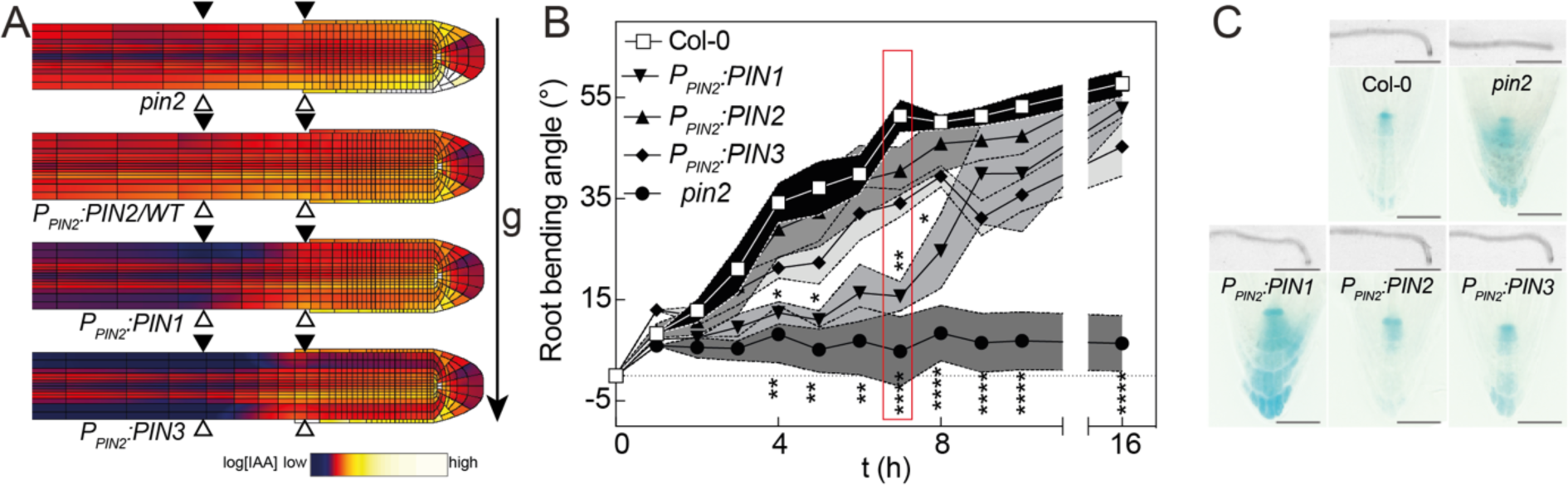
Response of PIN1, PIN2, PIN3 to a gravitropic stimulus. **(A)** Model auxin distribution patterns for the genotypes indicated. Simulated auxin pattern for 8 times faster transport rates in the columella and 2 times faster transport rates in the vasculature, representing columella localized fastest transporting PIN3 and vasculature localized intermediate fast transporting PIN1. The elongation zone is indicated by black arrowheads (upper side) and white arrowheads (lower side). The gravity vector is indicated. **(B)** Root bending kinetics of Col-0, *pin2*, *P_PIN2_:PIN1*, *P_PIN2_:PIN2* and *P_PIN2_:PIN3* on 0.5 MS medium. Red box highlights the 7 h time point. Data points are mean and SE of n = 3 (for *P_PIN2_:PIN1* and *P_PIN2_:PIN3)* or n = 6 (for *P_PIN2_:PIN2*) independent replicates consisting of 18-20 roots per genotype **(C)** Upper panel: Representative roots of Col-0, the *pin2* mutant, *P_PIN2_:PIN1*, *P_PIN2_:PIN2* and *P_PIN2_:PIN3* in *pin2* 7 h after the gravitropic stimulus on 0.5 MS medium. Scale bars represent 1 mm. Lower panel: Root tips of the respective genotypes crossed with auxin response reporter *Pro_DR5rev_:GUS*, stained 7 h after the gravitropic stimulus. Scale bars represent 50 µm. Groups were compared by one-way ANOVA, followed by Dunnett posthoc test with Col-0 as control group. Significant differences are indicated by “*”. n.s. not significant p-value ≥ 0.05, * p-value 0.01 to 0.05, ** p-value 0.001 to 0.01, *** p-value 0.0001 to 0.001, **** p-value < 0.0001.

### Probing the contribution of the loop to the transport properties of PINs by generating a “pseudocanonical” PIN

The data we obtained so far prompted us to investigate if features found in the loop can also be transferred to a non-canonical PIN. PIN8 is constitutively active and possesses a short loop but is closely related to PIN3, PIN4 and PIN7, suggesting that PIN8 lost the loop during evolution ^16,52^. We therefore introduced the PIN2 or the PIN3 loop between amino acid position 163 and 164 of PIN8 and investigated the biochemical and physiological features of these chimeras.

We found that the PIN8-2-8 and PIN8-3-8 chimeras were functional in the Xenopus oocyte assay and mediated NPA-sensitive IAA export, indicating that the chimaeras display all expected properties of PINs (Fig. 5A and Suppl. Fig. 7A) ^16,19^. Interestingly, we found that the PIN8-2-8 chimera, like PIN8, displayed constitutive IAA transport in the absence of a kinase whereas the PIN8-3-8 chimera, like a “typical” canonical PIN does not transport IAA without kinase activation (Fig. 5A). This clearly indicates that the loop is the decisive factor for PIN regulation. In both cases, the introduction of the loop led to an increase of the transport rate upon co-expression with a kinase. Surprisingly, the PIN8-2-8 chimera, upon co-expression of the PID kinase mediated IAA export that was significantly higher than either PIN2 or PIN8 and was similar to PIN3 (Fig. 5A and B compared to Fig 1D and G and Suppl-Fig. 1E and F). We next investigated the transport rates of PIN8 and the PIN8-2-8 chimera as a function of substrate concentration (Fig. 5B). We were able to fit a Michaelis-Menten model to PIN8 with a *K_m_* of 40 µM (Supplemental Table Statistics). This is in line with the *K_d_* established for this transporter and further supports the hypothesis that PINs possess substrate affinities that are much higher than the physiological IAA concentrations ^16^. As it was the case for the canonical PINs, we found that for the PID-activated PIN8-2-8 chimera transport rates increased and we were unable to fit a Michaelis-Menten model indicating that the chimera displays a lower affinity and the transport rates are in the seemingly linear range, far below *K_m_*. The PIN8-3-8 chimera exhibited transport properties of a typical canonical PIN: It showed no transport in the absence of a kinase and it could be activated by kinase co-expression so all features of the PIN8-3-8 chimera were almost identical to those of PIN3 (Fig. 5A). The data further support the conclusion that the loop together with the transmembrane context contributes to transport. We hypothesize that the activated loop, possibly due to the increased flexibility between the scaffold and the transport domain, increases the transport rate by promoting the flip-back from the outward open to the inward open configuration, the rate limiting step of the transport cycle ^16^. The observation that the PIN3 loop could autoinhibit the PIN8-3-8 chimera whereas the PIN2 loop in the PIN8-2-8 chimera was unable to do so is consistent with a previous study which suggests that the transmembrane domains and the loop coevolved and that PIN3 and PIN8 are more closely phylogenetically related ^16,34,35,52^.

**Figure 5.**
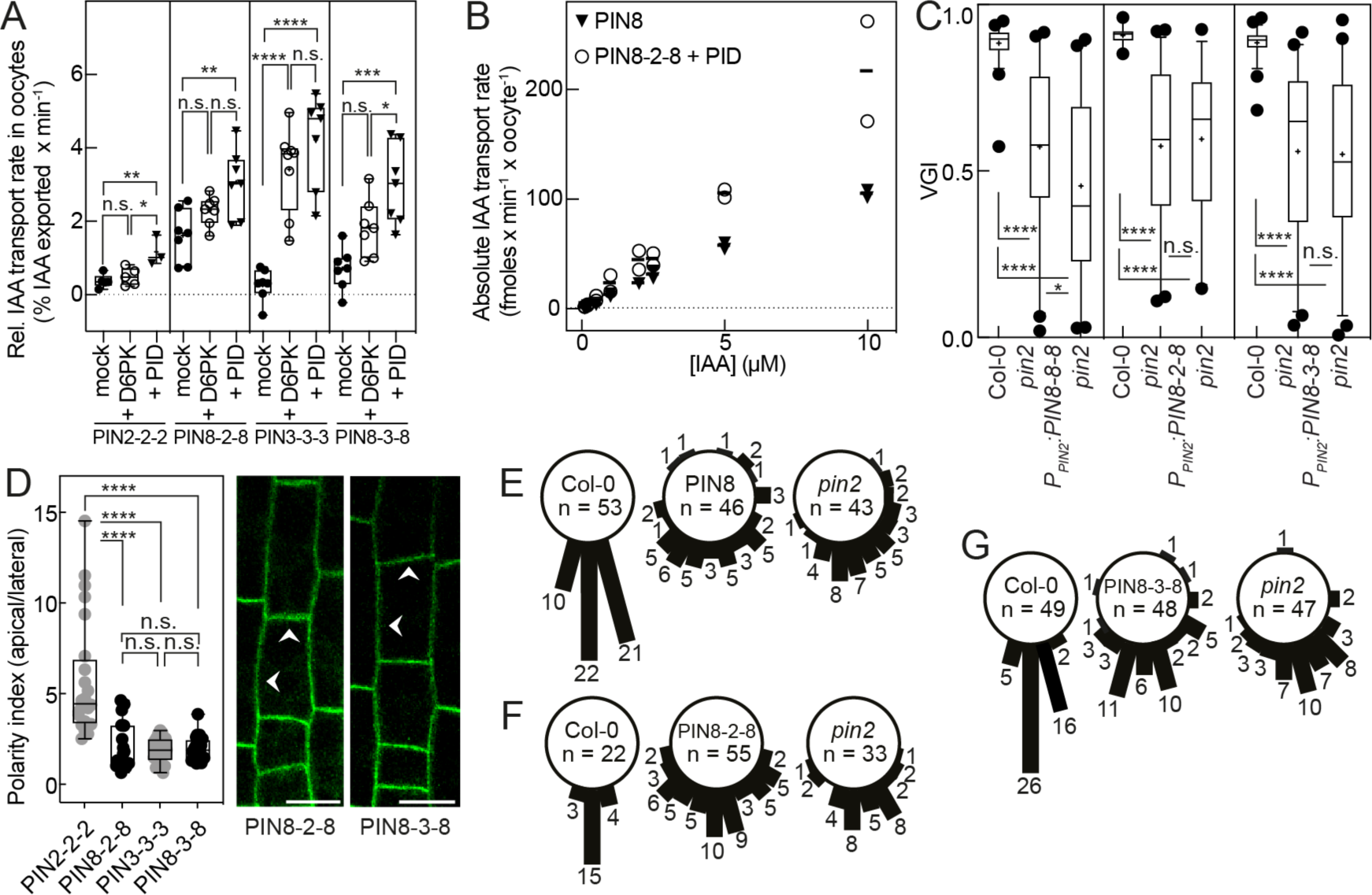
IAA transport properties, localization and potential to rescue the *pin2* phenotype by PIN8 and pseudocanonical chimeras. **(A)** Relative IAA transport rates of PIN2, PIN8-2-8, PIN3 and PIN8-3-8 expressed alone (●) or co-expressed with D6PK (○) or PINOID (▾) in oocytes. Data points are biological replicates using oocytes from different females (n = 3 for PIN2 + PINOID, n = 5 for PIN2/+ D6PK, n = 7 for PIN8-2-8/+ D6PK/+ PINOID and PIN8-3-8/+ D6PK/+ PINOID and PIN3/+ PINOID, n = 8 for PIN3 + D6PK). Groups were compared by one-way ANOVA, followed by Tukey’s post hoc test. PIN2 vs. PIN2 + D6PK: n.s., not significant, p-value 0.7584; PIN2 vs. PIN2 + PINOID: ** p-value 0.0056; PIN2 + D6PK vs. PIN2 + PINOID: * p-value 0.0152; PIN8-2-8 vs. PIN8-2-8 + D6PK: n.s. p-value 0.2076; PIN8-2-8 vs. PIN8-2-8 + PINOID: ** p-value 0.0036; PIN8-2-8 + D6PK vs. PIN8-2-8 + PINOID: n.s. p-value 0.1346; PIN3 vs. PIN3 + D6PK: **** p-value <0.0001; PIN3 vs. PIN3 + PINOID: **** p-value <0.0001; PIN3 + D6PK vs. PIN3 + PINOID: n.s. 0.3150; PIN8-3-8 vs. PIN8-3-8 + D6PK: n.s. 0.0534; PIN8-3-8 vs. PIN8-3-8 + PINOID: *** p-value 0.0002; PIN8-3-8 + D6PK vs. PIN8-3-8 + PINOID: * 0.0479. **(B)** Transport rates as function of [IAA]_in_. Data are mean and SE of n = 2 biological replicates using oocytes from different females. **(C)** VGIs of a representative homozygous T3 line of *P_PIN2_:PIN8*, *P_PIN2_:PIN8-2-8,* and *P_PIN2_:PIN8-3-8* in *pin2* (n = 46 for PIN8, n = 55 for PIN8-2-8, n = 48 for PIN8-3-8) in comparison to wildtype (n = 53 for PIN8, n = 22 for PIN8-2-8, n = 49 for PIN8-3-8) and mutant (n = 43 for PIN8, n = 33 for PIN8-2-8, n = 47 for PIN8-3-8). Box plots range from 25th to 75th percentiles, whiskers mark the 5th and 95th percentiles and the median is indicated. Points below and above the whiskers are drawn as individual points. The mean is indicated by (+). Groups were compared by one-way ANOVA followed by Tukey’s posthoc test. n.s. not significant p-value 0.8804 (PIN8-2-8), 0.9751 (PIN8-3-8), * p-value 0.0225, **** p-value <0.0001. **(D)** Polarity indices of PIN8-2-8-GFP and PIN8-3-8-GFP expressed from a *PIN2* promoter fragment in epidermal cells at the onset of elongation in *pin2*. PIN2 and PIN3 were included for comparison as presented in Figure 2. Mean intensities of ROIs at the apical and the lateral side of the cells were used to calculate the ratios. The arrows mark predominant localization. Scale bars represent 10 µm. Box plots range from 25th to 75th percentiles, whiskers mark the minimum and maximum values and the median is indicated. Groups were compared by a one-way ANOVA, followed by Tukey’s posthoc test. **** p-value <0.0001, n.s. not significant p-value 0.9749 (PIN8-2-8 vs. PIN3), 0.9953 (PIN8-2-8 vs. PIN8-3-8), 0.9978 (PIN8-3-8 vs. PIN3). n = 24 cells per genotype. **(E-G)** Root angles between root tip and gravity vector of a representative homozygous T3 line of *P_PIN2_:PIN8, P_PIN2_:PIN8-2-8* and *P_PIN2_:PIN8-3-8* in the *pin2* mutant background in comparison to wildtype and mutant. Numbers are individual seedlings.

We were interested to see if the pseudocanonical PINs also can complement the VGI of the *pin2* mutant. To this end we again expressed the chimeras under control of the *PIN2* promoter fragment and scored the VGI of a representative T3 identified among 20 independent lines from a segregating T2 population. We found that the chimeras localized non-polarly to the plasma membrane and the chimeras, as well as PIN8, failed to complement all aspects of the mutant phenotype (Fig. 5D and Suppl. Fig. 7B-F). Additionally, the PIN8-2-8 chimera, despite containing the PIN2 loop and showing apical localization, was unable to respond to the gravitropic stimulus (Supplemental Figure 7E-G). This once more indicates that the interplay of transmembrane domain and loop *in plant*a is more complex and goes beyond localization to mediate the flux rates required for proper physiological function. In summary we could show that PIN8-3-8 behaves like a canonical PIN whereas PIN8-2-8 displays only some of these features. It will be interesting to identify the regions in the loop that interact with the transmembrane domains to autoinhibit the transporter.

## Conclusions

The data presented here demonstrate that PINs differ significantly in their transport properties. Our data also show that these transport rates, and not merely tissue specific PIN polarity, play a major role in their physiological function. Finally, we demonstrate that both loop and transmembrane domains of the PIN proteins contribute to their localization and transport properties, and that this “domain collaboration” has arisen through co-evolution. How this collaboration works on a mechanistic level and whether the subtle kinase-dependent differences we observed in the transport assays have a role in this process that translates into physiological differences remains to be answered in future studies.

## Supporting information

Janacek et al Extended Data Figures

Janacek Extended Data Table 1

## Acknowledgements

This work was funded by DFG3468/6-1, DFG3468/6-3 and SFB924 to UZH. We thank Angela Alkofer and Helene Prunkl for excellent technical assistance and Xenopus maintenance.

## Methods

### Cloning procedure

Plant expression vectors were generated by a modified GreenGate cloning protocol ^53^. The BsaI-recognition site and, if necessary, suitable overhangs were added to the DNA fragments by PCR. The DNA fragments of interest were amplified from genomic DNA or synthesized by Geneart, Thermo Fisher (Regensburg, Germany). The transmembrane and loop domains of PIN1, PIN2, PIN3 and PIN8 were defined according to ^45^. PIN1: TM1 1–156 aa, loop 157–459 aa, TM2 460–623 aa. PIN2: TM1 1–156 aa, loop 157–484 aa, TM2 485–623 aa. PIN3: TM1 1–156 aa, loop 157–477 aa, TM2 478–641 aa. PIN8: TM1 1–163 aa, loop 164–204 aa, TM2 205–367 aa. The CDS of eGFP was inserted C-terminally of amino acid positions 432 (PIN1), 301 (PIN2) and 451 (PIN3) or at equivalent positions in the loop domain in the PIN chimeras ^45^. The loops of PIN1, PIN2 or PIN3 were inserted C-terminally of position 163 in PIN8.

### Oocyte Transport assay

For cloning into pOO2 ^54^, the coding sequences were amplified with 5’-phosphorylated oligos and purified by gel electrophoreses and sequenced. The efflux experiments were performed as described ^55^. For cRNA synthesis, the mMESSAGE mMACHINE™ SP6 Transcription kit and the MEGAclear Purification of Transcription Reactions kit from Thermo Fisher Scientific were used according to the manufacturer’s protocol. Oocytes were injected with 150 ng/µl PIN cRNA and 75 ng/µl cRNA of respective kinase. ^3^H-IAA (25 Ci/mmol, 1 mCi/mmol) was purchased by ARC (Saint Louis, USA) or RC Tritec (Teufen, Switzerland). After four days, oocytes were injected with substrate to reach the desired internal IAA concentration. For each ach time point (0, 5, 10, 15 min for PIN3, PIN2-3-2, PIN3-1-3, PIN3-2-3, PIN8 or 0, 7.5, 15, 30 min for PIN1, PIN2, PIN1-2-1, PIN2-1-2, PIN8-2-8, PIN8-3-8) at least seven oocytes were used. The transport rates for each concentration were calculated by linear regression. Experiments were performed at least three times with oocytes from different *X. laevis* females.

### (Phospho)proteomic sample preparation

Oocytes (n = 50) expressing either PIN1, PIN2, PIN3 alone or PIN co-expressed with YFP-D6PK or PINOID were collected without oocyte buffer in a reaction tube (2 ml). The oocytes were homogenized in 2 ml Lysis buffer (Tris-HCl pH 8.0 50 mM, ½ tablet PhosSTOP (Roche), 1x cOmplete^TM^ EDTA-free proteinase inhibitor cocktail (Sigma-Aldrich), 1 % SDC) and centrifuged (2000 *g*, 10 min, 4 °C). The supernatant without yolk was transferred to a reaction tube suitable for ultracentrifugation and centrifuged (150 000 *g*, 30 min, 4 °C). The cytosolic fraction (supernatant) and the membrane fraction (pellet) were split and the membrane fraction was resuspended in 400 µl Lysis buffer. All samples were stored at –80 °C before preparation for LC-MS/MS analysis. Proteins were precipitated over night with 10% tricholoroacetic acid in acetone at - 20° C and subsequently washed two times with ice-cold acetone. Dry samples were incubated with urea digestion buffer (8 M urea, 50 mM Tris-HCl pH 8.5, 1 mM DTT, cOmplete^TM^ EDTA-free protease inhibitor cocktail (PIC, Roche, Basel, Switzerland), Phosphatase inhibitor (PI-III; in-house, composition resembling Phosphatase inhibitor cocktail 1,2 and 3 from Sigma-Aldrich, St. Louis, USA) for 1 h. Protein concentration was determined with a Bradford assay ^56^. For each sample 20 µg (cytoplasm) or 80 µg (membrane) of protein was reduced (10 mM DTT), alkylated (55 mM chloroacetamide), and diluted 1:5 with digestion buffer (50 mM Tris-HCl pH 8.5, 1 mM CaCl_2_). In-solution digestion with trypsin (1:100 w/w) at 37°C was performed in two steps (4h; overnight). Digested samples were acidified with formic acid (FA) and centrifuged at 14,000 *g* for 15 min at 4°C. Cytoplasm samples were desalted on self-packed StageTips (three disks, Ø 1.5 mm C18 material, 3M Empore^TM^, elution solvent 0.1% FA in 50% ACN) and the peptides dried in a vacuum concentrator prior to liquid chromatography-coupled mass spectrometry (LC-MS) analysis. For the membrane samples Fe^3+^-IMAC was performed as described previously with some adjustments ^57^. SepPac desalted peptide samples were re-suspended in ice-cold IMAC loading buffer (0.1% TFA, 40% acetonitrile). For quality control, 1.5 nmol of a synthetic library of phosphopeptides and their corresponding non-phosphorylated counterpart sequence (B2 and F1) ^58^ were spiked into each sample prior to loading onto a Fe^3+^-IMAC column (Propac IMAC-10 4×50 mm, Thermo Fisher Scientific). The enrichment was performed with Buffer A (0.07% TFA, 30% acetonitrile) as wash buffer and Buffer B (0.315% NH_4_OH) as elution buffer. Collected full proteome and phosphopeptide fractions were vacuum-dried, reconstituted in 0.1% FA or 0.1% Fa, 50 mM Citrate, respectively, desalted on self-packed stage tips and dried down prior to LC-MS analysis.

### LC-MS/MS measurement

Dry peptides were re-suspended in 0.1% FA in HPLC grade water and spiked with PROCAL retention time standard peptide ^59^. LC-MS/MS analysis was performed on a Q Exactive HF Orbitrap (Thermo Fisher Scientific) coupled on-line to a Dionex Ultimate 3000 RSLCnano system. The liquid chromatography setup consisted of a 75 μm x 2 cm trap column and a 75 μm x 40 cm analytical column, packed in-house with Reprosil-Pur C18 ODS-3 5 μm or Reprosil Gold C18 3 μm resin (Dr. Maisch GmbH), respectively. Peptides were loaded onto the trap column using 0.1% FA in water at a flow rate of 5 μL/min and separated using a 50 min linear gradient from 4% to 32% of solvent B (0.1% (v/v) formic acid, 5% (v/v) DMSO in acetonitrile) at 300 nL/min flow rate. nanoLC solvent A was 0.1% (v/v) formic acid, 5% (v/v) DMSO in HPLC grade water ^60^. A 50 min two step gradient from 4% to 27% solvent B was used for the phosphoproteome samples.

Peptides were ionized using a spray voltage of 2.2 kV and a capillary temperature of 275°C. The instrument was operated in data-dependent mode, automatically switching between MS1 and MS2 scans. Full scan MS1 spectra (m/z 360 – 1300) were acquired with a maximum injection time of 10 msec at 60,000 resolution and an automatic gain control (AGC) target value of 3e6 charges. For the top 20 precursor ions, high resolution MS2 spectra were generated in the Orbitrap with a maximum injection time of 25 msec at 15,000 resolution (isolation window 1.7 m/z), an AGC target value of 1e5 and normalized collision energy of 25%. The underfill ratio was set to 1% with a dynamic exclusion of 20 sec. Only precursors with charge states between 2 and 6 were selected for fragmentation. For the phosphoproteome analysis, the top 15 MS2 spectra were acquired with a maximum injection time of 100 msec, an AGC target value of 2e5 and a dynamic exclusion of 25 s.

### (Phospho)proteomic data analysis

Thermo raw files for membrane full proteome and phosphoproteome samples were processed together as two separate parameter groups using MaxQuant software (v. 1.6.3.3) with standard settings unless otherwise described ^61^. MS/MS spectra were searched against Araport11 ^62^ protein coding genes (Araport11_genes.201606.pep.fasta; download 06/2016), the Xenopus reference proteome (UP00186698; uniprot download 05/2018) and spike-in phosphopeptide and PROCAL peptide library sequences ^58,59^, with trypsin as protease and up to two allowed missed cleavages. Carbamidomethylation of cysteines was set as fixed modification and oxidation of methionine and N-terminal acetylation as variable modifications. For the phosphoproteome parameter group phosphorylation of serine, threonine or tyrosine was added as variable modification. Results were filtered to 1% PSM, protein and Site FDR. Raw data files for the cytoplasm proteome were processed in a separate MaxQuant search using the same settings as for the membrane full proteome.

For protein abundance comparisons PIN1 (AT1G73590), PIN2 (AT5G57090), PIN3 (AT1G70940), D6PK (AT5G55910) and PINOID (AT2G34650) LFQ peptide intensities were summed up for the respective samples and length corrected with the number of iBAQ peptides. Phosphorylation site intensities were normalized with the total peptide intensity (allPeptides.txt) ratio of the respective sample and filtered for sites identified with two valid values per grouping. Statistic comparison of D6PK and PINOID phosphorylation was performed in Perseus using Student T-test with standard settings (missing values imputed from normal distribution, 1.8 downshift, 0.3 width, separately for each column; s0=0; Permutation based FDR 0.05) ^63^.

The mass spectrometry proteomics data have been deposited to the ProteomeXchange Consortium via the PRIDE partner repository with the dataset identifier PXD044850 (user name, password: 3xBdgLSX) ^64^.

### Plant growth conditions

For growth in pots, the seeds were spread on moist soil and incubated in the growth chamber with a cover under long day conditions (16 h light, 8 h dark, 21-23 °C). After one week, the cover was removed and the plants were separated to individual pots, if necessary. For growth on plates, the seeds were sterilized with chlorine gas (6 %(v/v), 1 h) and homogenously spread on the agar or placed individually with an autoclaved toothpick. Plates were sealed and incubated at 4 °C in the dark for two days. Afterwards the plates were placed vertically in plant growth chambers (Sanyo or Panasonic) under long day conditions (16 h light, 8 h dark, 21 °C).

Solid 0.5 MS medium contained 2.15 g/l Murashige & Skoog medium, including B5 vitamins incl. Gamborg B5 vitamins (Duchefa Farma, Harleem, Netherlands), 0.5 g/l MES monohydrate (Carl Roth, Karlsruhe, Germany) and 0.8 % (w/v) plant agar (Duchefa Farma). Solid Growth medium contained 4.3 g/l MS medium, including B5 vitamins incl. Gamborg B5 vitamins, 0.5 g/l MES monohydrate, 0.7 % (w/v) plant agar and 1 % (w/v) saccharose.

### Plant transformation and genotyping

Plants were transformed using *Agrobacterium tumefaciens* GV3101 according to ^65^. In order to select for positive transformants, the seeds were sprayed with a Basta solution (1 % v/v). Seedlings that survived the treatment were transferred to single pots and genotyped by PCR.

### Immunocytochemical techniques

Localization of proteins by immunocytochemical techniques was performed as described ^66,67^. Antibodies were used as published previously ^68^.

### Modeling of IAA distribution in roots

Auxin dynamics were modeled on a two-dimensional cross section of an idealized *Arabidopsis thaliana* root tip anatomy containing at its distal end a quiescent center, surrounded by stem cell niche, columella, and root cap, and shootward from the QC consisting from inside to outside epidermal, cortical, endodermal, pericycle and multiple vascular tissue cell files. The model incorporates experimentally derived cell type and root zone specific patterns of the AUX/LAX auxin importers, the polarly localized PIN auxin exporters, and in addition to baseline cell level auxin production and degradation elevated levels of auxin production around the QC, in the columella and lateral root cap, as done previously ^5,26,69^. Under standard model settings ^5^ PIN efflux rates are equal across all cell types and hence PIN types. Taking PIN2 transport rates as a default, we modeled the faster PIN3 mediated auxin transport by elevating columellar PIN efflux rates 8-fold, whereas to model the 2-fold higher transport rate of PIN1 we elevated vascular and pericycle PIN efflux rates 2-fold. Epidermal, cortical and endodermal PIN efflux rates were kept identical to standard settings.

In simulations of gravitropic stimulation, columellar PIN3 patterns were polarized to simulate their reorientation in response to statolith sedimentation ^70^. In contrast to the default apolar PIN pattern in these cells, downward oriented membrane faces received 1.5 times more PINs whereas all other membrane faces received 0.3-fold lower PIN levels.

To simulate the *pin2* mutant, PIN levels in lateral root cap, epidermis and cortex were reduced to 0.1-fold their original value, reflecting that in *pin2* mutants there is a partial takeover by PIN1 becoming active in the PIN2 domain with correct polarity yet insufficiently to restore gravitropic response ^7^. To simulate *PIN2* promotor mediated *PIN1* expression in *pin2* mutants, we replaced the polar PIN2 pattern with an apolar PIN1 pattern in epidermis and cortex. To simulate *PIN3* expression in the *PIN2* domain in *pin2* mutants, we replaced the polar PIN2 pattern with an apolar PIN3 pattern in epidermis and cortex and increased local PIN transport rates 8-fold.

The model is grid based, meaning individual cells and cell walls consist of a collection of grid points, and auxin dynamics are modeled as a partial differential equation. Grid based auxin dynamics were solved using an Alternating Direction Implicit (ADI) integration scheme, using a time step of 0.2s and a space step of 2microm, again as was done before ^5,26,69^

### Gravitropism assays

Sterilized seeds of Col-0, *pin2* mutant and the PIN T-DNA line were plated in two sets of 10 seeds per genotype on the plate containing 0.5 MS + 1 % sucrose. In order to minimize plate effects, the position of the genotypes was rotated on different plates, resulting in six plates and 120 seeds for each genotype per investigated PIN rescue line. The plates were sealed and incubated in the dark at 4 °C for two days. Afterwards the plates were placed vertically into a plant growth chamber and scanned 5 days later. The root angle between the root origin and the root tip was measured for every seedling, using the SNT plugin of FIJI and a script to give the value of the root angle simultaneously ^71,72^. For each PIN construct, 5-10 individual segregating T2 lines were analyzed as described before. One representative line was propagated to the next generation, in order to generate a homozygous line. The representative homozygous PIN T-DNA line was plated with wildtype and mutant seedlings on the same plates with only one plate per plate layout (n = 60 seeds). The plates were processed as described earlier. The VGI of the homozygous lines was calculated using FIJI ^33^.

### Root bending assay

The homozygous PIN T-DNA lines of interest, the wildtype Col-0 and the *pin2* mutant were grown on plates containing the indicated medium for five days, after two days of stratification at 4 °C in the dark. Two times five seedlings were transferred to a new plate containing medium as indicated (either 0.5 MS + 1 % sucrose, or 0.5 MS without vitamins) and the root was straightened. The plate was turned 90 ° and was placed into a growth chamber (Sanyo, Moriguchi, Japan), together with an IR LED light module. A Raspberry Pi (Raspberry Pi Foundation, UK) equipped with an IR camera was placed in front of the plate and an image was taken every five minutes. The angle between the root body and the tip was measured every hour from 1-10 hours after turning and after 16 h.

### GUS staining

The seeds of homozygous PIN T-DNA lines, the wildtype Col-0 and the *pin2* mutant were grown on plates containing 0.5 MS + 1 % sucrose for five days, after two days of stratification at 4 °C in the dark. The whole seedlings were transferred to a 6-well plate containing GUS staining solution (100 mM NaPO_4_ pH 7.0, 100 mM EDTA pH 7.0, 1 mM K₃[Fe(CN)₆], 1 mM K₄[Fe(CN)₆]·3H₂O, 0.1 % Triton X-100 in H_2_O, 0.5 mg/ml X-Gluc in DMF) and incubated for 1 h at 37 °C with the plate covered in aluminum foil. Afterwards the seedlings were washed three times in buffer (50 mM NaPO_4_ pH 7.0). The roots were mounted in chloral hydrate solution (50 % (w/v) chloral hydrate, 10 % (v/v) glycerol) and imaged at an Olympus BX61 Upright microscope (Hamburg, Germany).

### Microscopy and signal quantification

In order to image the PIN localization in Arabidopsis roots or to image the R2D2 auxin reporter, an Olympus BX61 microscope with a FV1000 confocal laser scanning unit (Olympus, Hamburg, Germany) or a Leica TCS SP8 confocal microscope (Leica, Wetzlar, Germany) were used. The brightness and contrast were adjusted in all images and the scale bar was automatically included using FIJI. All measurements for the polarity index or the R2D2 signal analysis were performed in FIJI.

The polarity index was determined by calculating the ratio of the GFP signal at the apical and lateral PM of root epidermal cells. A square (3 px x 15 px) was defined as region of interest (ROI) and four cells from two roots of three independent segregating lines were analyzed per genotype.

In order to analyze the R2D2 signal ^39^, a maximum projection of 3-8 images with 2.0 µm intervals of either the epidermal or cortical cell file was used. A round ROI covered the nucleus. The R2 (mDII signal) to D2 (DII signal) ratio was calculated of the first five cells after the anticlinal division of the epidermis/lateral root cap initial cell (Q) and five cells at the transition zone (T). The GUS-stained roots were imaged at an Olympus BX61 Upright microscope (Hamburg, Germany).

### Statistics

All data were plotted with GraphPad Prism V9 (Boston, USA). Statistical tests as indicated were performed using the default settings of GraphPad Prism V9.

