## Supplementary material for "Transport properties of canonical PIN-FORMED proteins and the role of the loop domain in auxin transport": Janacek et al Extended Data Figures

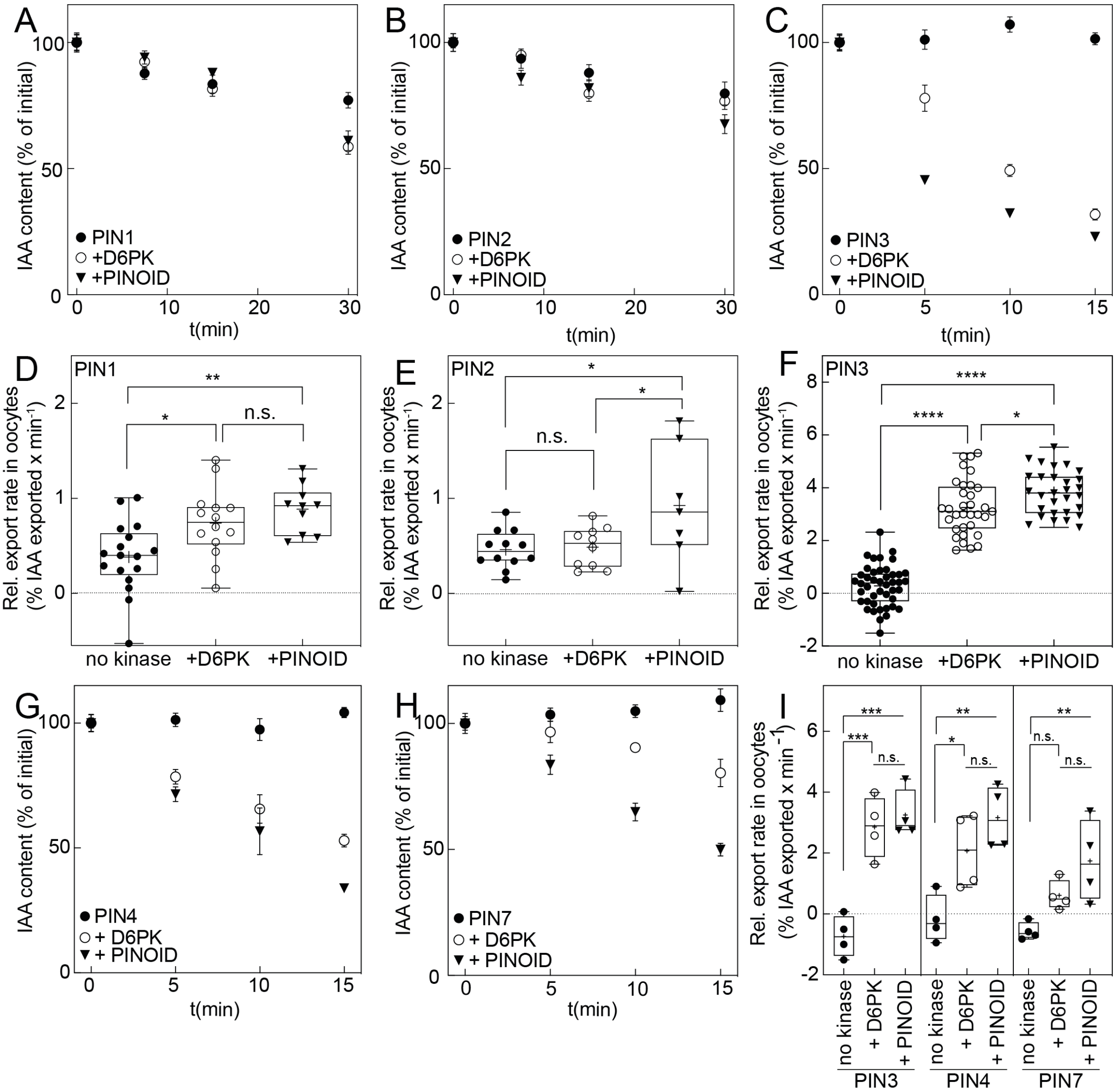

**Supplemental Figure 1. IAA transport properties of canonical PINs in Xenopus oocytes. (A)** Representative time-course experiments for PIN1 expressed without (●) with D6PK (○) or with PID (▼). Time points are mean and SE of (n = 8-10 individual oocytes). **(B)** Representative time-course experiments for PIN2 expressed without (●) with D6PK (○) or with PID (▼). Time points are mean and SE of (n = 8-10 individual oocytes). **(C)** Representative time-course experiments for PIN3 expressed without (●) with D6PK (○) or with PID (▼). Time points are mean and SE of (n = 8-10 individual oocytes). **(D)** Relative IAA export rates for PIN1 (n = 10-17 biological replicates) at $[IAA_{in}]$ = 1 µM. **(E)** Relative IAA export rates for PIN2 (n = 7-12 biological replicates) at $[IAA_{in}]$ = 1 µM. **(F)** Relative IAA export rates for PIN3 (n = 30-44 biological replicates) at $[IAA_{in}]$ = 1 µM+. Box plots range from the 25th to 75th percentile and the median is shown, the mean is represented by (+). Groups were compared by one-way ANOVA, followed by Tukey's post-hoc test (PIN1 vs. PIN1 + D6PK: * p-value 0.0178, PIN1 vs. PIN1 + PINOID: ** p-value 0.0021, PIN1 + D6PK vs. PIN1 + PINOID: n.s., not significant, p-value 0.5589; PIN2 vs. PIN2 + D6PK: n.s. p-value 0.9838, PIN2 vs. PIN2 + PINOID: * p-value 0.0265, PIN2 + D6PK vs. PIN2 + PINOID: * p-value 0.0458; PIN3 vs. PIN3 + D6PK: **** p-value <0.0001, PIN3 vs. PIN3 + PINOID: **** p-value <0.0001, PIN3 + D6PK vs. PIN3 + PINOID: * p-value 0.0341). **(G)** Representative time-course experiments for PIN4 expressed without (●) with D6PK (○) or with PID (▼). Time points are mean and SE of (n = 8-10 individual oocytes). **(H)** Representative time-course experiments for PIN7 expressed without (●) with D6PK (○) or with PID (▼). Time points are mean and SE of (n = 8-10 individual oocytes). **(I)** Relative IAA export rates at [1 µM] $[IAA_{in}]$ = 1 µM for PIN3, PIN4 and PIN7. n = 4 biological replicates for all constructs. Box plots range from the minimum to maximum percentile and the median is shown, the mean is represented by (+). Groups were compared by one-way ANOVA, followed by Tukey's post-hoc test (PIN3 vs. PIN3 + D6PK: *** p-value 0.0005, PIN3 vs. PIN3 + PINOID: *** p-value 0.0002, PIN3 + D6PK vs. PIN3 + PINOID: n.s., not significant, p-value 0.7894, PIN4 vs. PIN4 + D6PK: * p-value 0.0338, PIN4 vs. PIN4 + PINOID: ** p-value 0.0037, PIN4 + D6PK vs. PIN4 + PINOID: n.s. p-value 0.3447, PIN7 vs. PIN7 + D6PK: n.s. p-value 0.1750, PIN7 vs. PIN7 + PINOID: ** p-value 0.0095, PIN7 + D6PK vs. PIN7 + PINOID: n.s. p-value 0.1931).

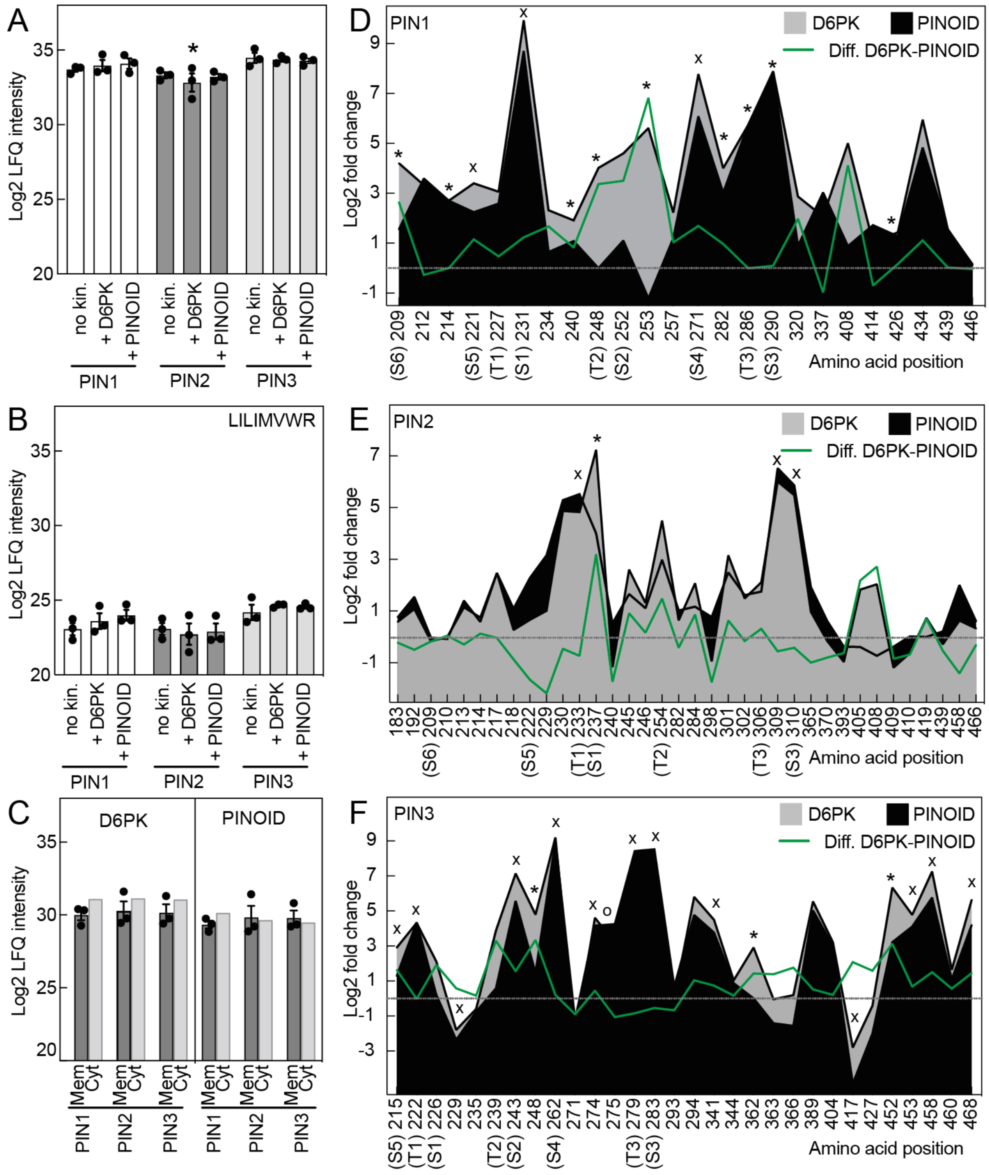

**Supplemental Figure 2. Proteome and phospho-proteome analyses of PINs and D6PK or PINOID in *Xenopus* oocytes. (A)** LFQ intensity of PIN1, PIN2 and PIN3 protein in the membrane fraction without kinase (n = 3) and with D6PK (n = 3) or PINOID (n = 3). Data are mean and SE. Data analyzed in their respective group (PIN vs. + D6PK vs. + PINOID) by one-way ANOVA, followed by Tukey's posthoc test showed no significant difference (PIN1 vs. PIN1 + D6PK n.s. 0.8239, PIN1 vs. PIN1 + PINOID n.s. 0.6612, PIN1 + D6PK vs. PIN1 + PINOID n.s. 0.9535. PIN2 vs. PIN2 + D6PK n.s. 0.6559, PIN2 vs. PIN2 + PINOID n.s. 0.9809, PIN2 + D6PK vs. PIN2 + PINOID n.s. 0.7614. PIN3 vs. PIN3 + D6PK n.s. 0.9442, PIN3 vs. PIN3 + PINOID n.s. 0.8343, PIN3 + D6PK vs. PIN3 + PINOID n.s. 0.9642). **(B)** LFQ intensity of the peptide LILIMVWR shared between the three PINs in the membrane fraction (n = 3). Data represent mean and SE. Individual groups (PIN vs. + D6PK vs. + PINOID) were compared by one-way ANOVA, followed by Tukey's post hoc test. ns in Figure **(C)** LFQ intensity of D6PK and PINOID in the membrane fraction (n = 3) and the cytosolic fraction (n = 1) of oocytes expressing PIN and the kinase indicated. Data represent mean and SE. **(D-F)** Phosphorylated sites in PIN1, PIN2 and PIN3 in comparison to PIN without kinase. The green line indicates the difference between PIN with D6PK or PINOID. (X) marks sites phosphorylation sites that are significantly ($p < 0.05$) increased in response to both kinases, (*) marks phosphorylation sites that are significantly ($p < 0.05$) increased in response to D6PK and (o) marks phosphorylation sites that are significantly ($p < 0.05$) increased in response to PINOID (Student's t-test two sided unpaired, with multiple-testing correction Permutation-based FDR 0.05).

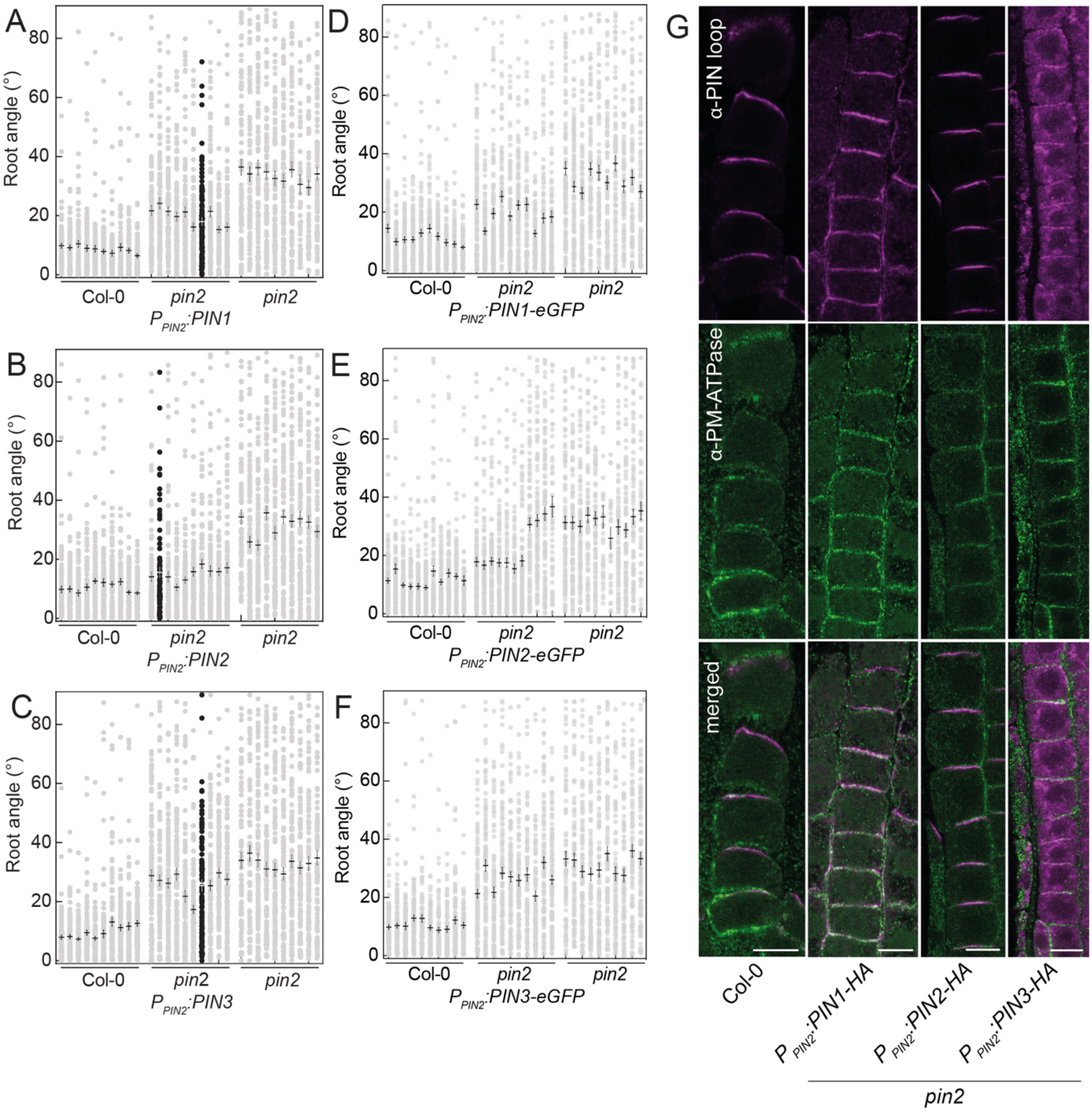


**Supplemental Figure 3. Identification of representative lines of canonical PINs with respect to complement the *pin2* mutant phenotype in a segregating T2 population. (A)** *$P_{PIN2}$:PIN1*, **(B)** *$P_{PIN2}$:PIN2*, **(C)** *$P_{PIN2}$:PIN3*, **(D)** *$P_{PIN2}$:PIN1-eGFP*, **(E)** *$P_{PIN2}$:PIN2-eGFP* **(F)** *$P_{PIN2}$:PIN3-eGFP* in *pin2*. Data points are individual seedlings. Transgenic lines (99 < n < 120) in comparison to wildtype (98 < n < 120) and mutant (47 < n < 119). The mean and SE are indicated. The representative line used for analysis of the T3 progeny is represented by black dots. **(G)** Immunolocalization of PIN1-HA, PIN2-HA and PIN3-HA at the transition zone as specified by cell elongation PIN loops was decorated with loop specific antisera (magenta), plasmamembrane ATPase is shown in green.

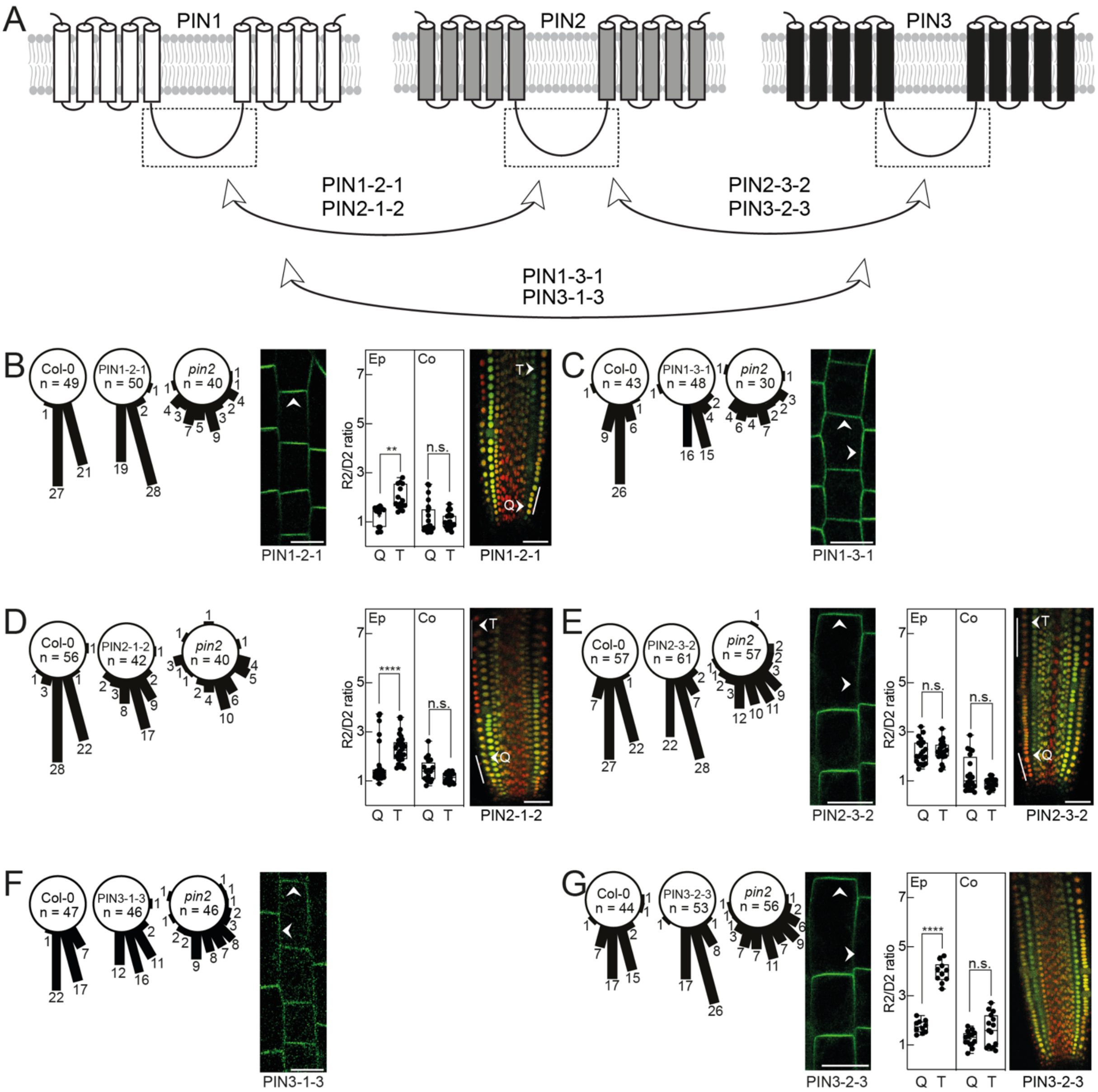

**Supplemental Figure 4. Relative transport rate of chimera between canonical PINs in *Xenopus* oocytes. (A)** Schematic representation of the domain swap approach. **(B)** $P_{PIN2}$*:PIN1-2-1*. **(C)** $P_{PIN2}$*:PIN1-3-1*. **(D)** $P_{PIN2}$*:PIN2-1-2*, **(E)** $P_{PIN2}$*:PIN2-3-2*, **(F)** $P_{PIN2}$*:PIN3-1-3*, **(G)** $P_{PIN2}$*:PIN3-2-3.* Root angles between root tip and gravity vector of a representative homozygous T3 line compared to wildtype and mutant. Numbers are individual seedlings. Localization of PIN1-2-1-eGFP, PIN1-3-1-eGFP, PIN2-3-2-eGFP, PIN3-1-3-eGFP and PIN3-2-3-eGFP in epidermal cells at the onset of elongation in *pin2*. The arrows mark predominant localization. Scale bars represent 10 µm. Ratio of mDII to DII signal in epidermis (Ep) and cortex (Co). The cells indicated by white lines are the first five epidermal cells after anticlinal division of the epidermal/LRC initial cell (Q) and five cells at the transition zone as specified by cell elongation (T) which were measured for individual roots (n = 15-30 for epidermis and n = 15-25 for cortex). The overlay of DII (green) and mDII (red) signal is shown. Scale bars represent 40 µm. Box plots show the 25th and 75th percentiles, whiskers mark the minimum and maximum values. The line in the box represents the median. Groups were compared by one-way ANOVA, followed by Tukey's post hoc test (PIN1-2-1: ** p-value 0.0077, n.s. not significant p-value 0.9991. PIN2-1-2: **** p-value <0.0001, n.s. p-value 0.4792. PIN2-3-2: n.s. p-value >0.9999 (Ep), n.s. p-value 0.1736 (Co). PIN3-2-3: **** p-value <0.0001, n.s. p-value 0.4814).

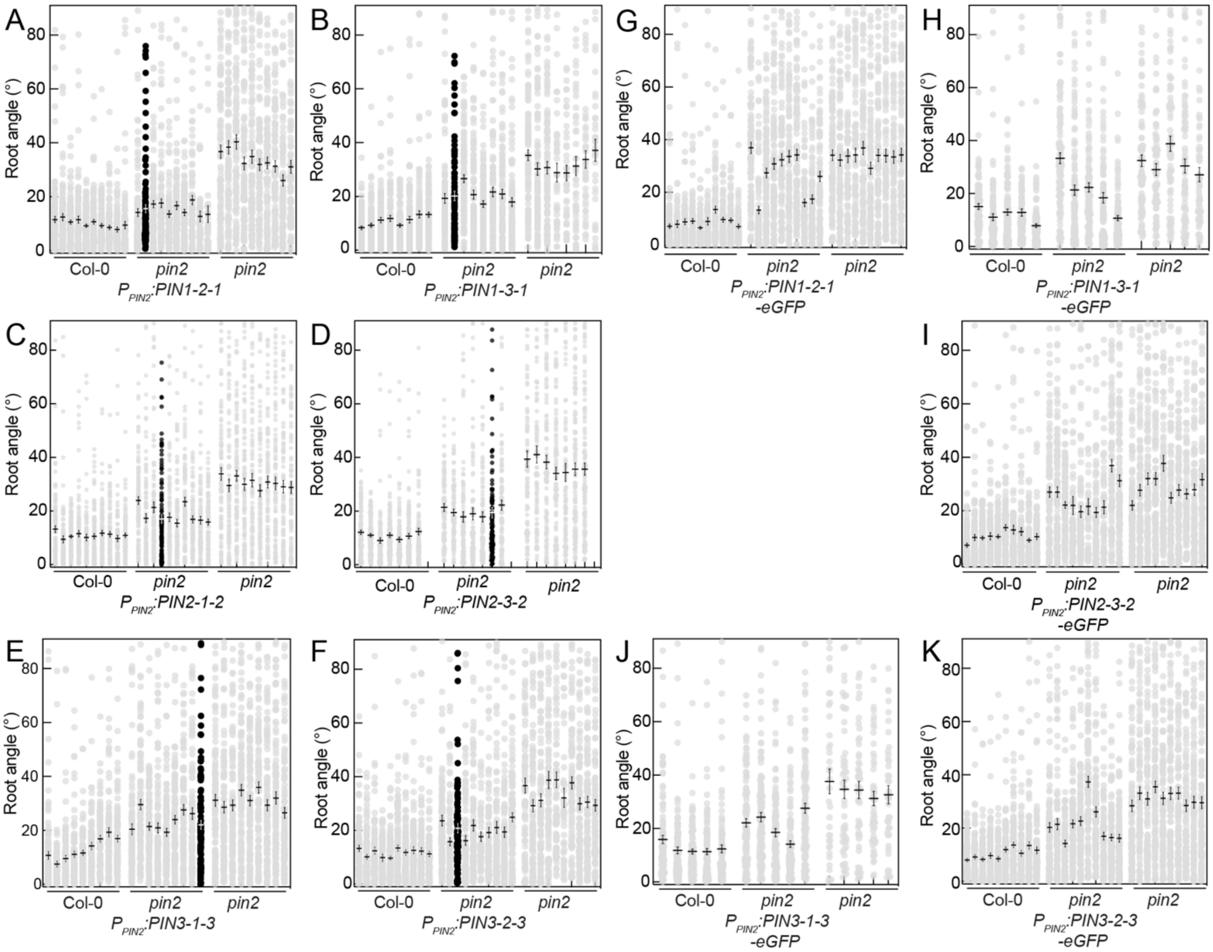


**Supplemental Figure 5. Identification of representative lines of chimeras between canonical PINs with respect to complement the *pin2* mutant phenotype in a segregating T2 population. (A)** $P_{PIN2}$:*PIN1-2-1*, **(B)** *Po*$_{PIN2}$:*PIN1-3-1*, **(C)** $P_{PIN2}$:*PIN2-1-2*, **(D)** $P_{PIN2}$:*PIN2-3-2*, **(E)** $P_{PIN2}$:*PIN3-1-3*, **(F)** $P_{PIN2}$:*PIN3-2-3* and **(G-K)** the corresponding eGFP-tagged variants in the *pin2* mutant background. Root angle of independent segregating T2 lines between root tip and gravity vector. Data points are individual seedlings of transgenic lines (37 < n < 120) compared to wildtype (97 < n < 120) and mutant (47 < n < 120). The mean and SE are indicated. The representative line used for analysis of the T3 progeny is represented by black dots in A-F.

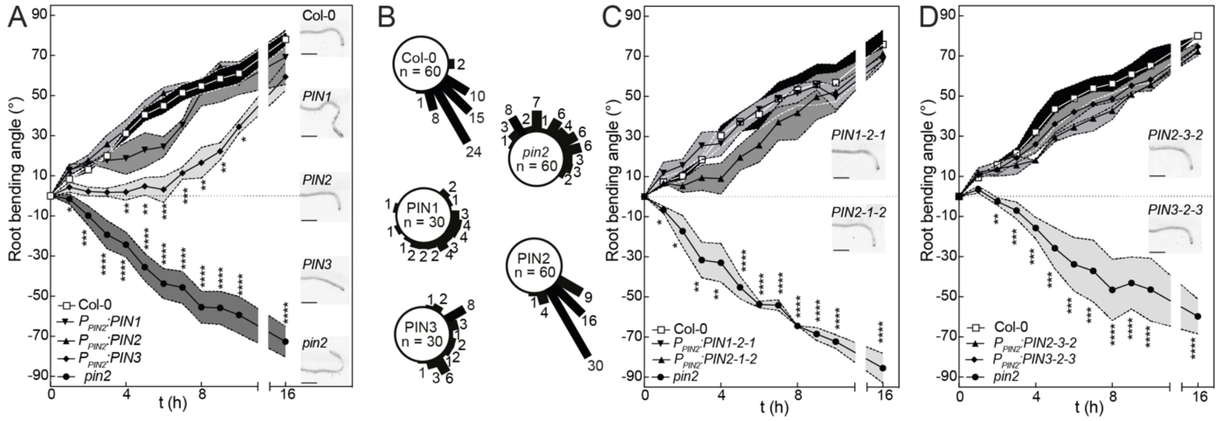


**Supplemental Figure 6. Response of the PIN chimeras to a gravitropic stimulus on plates with sucrose. (A)** Root bending kinetics of $P_{PIN2}$*:PIN1*, $P_{PIN2}$*:PIN2* and $P_{PIN2}$*:PIN3* in *pin2* compared to wildtype and mutant. Insets show representative roots of the genotypes indicated 7 h after the gravitropic stimulus. Data points are mean and SE of n = 3 (for $P_{PIN2}$*:PIN1* and $P_{PIN2}$*:PIN3)* or n = 6 (for $P_{PIN2}$*:PIN2*) independent replicates consisting of 18-20 roots per genotype. Groups were compared by one-way ANOVA, followed by Dunnett post hoc test with Col-0 as control group. Significant differences are indicated by "*". n.s. not significant p-value ≥ 0.05, * p-value 0.01 to 0.05, ** p-value 0.001 to 0.01, *** p-value 0.0001 to 0.001, **** p-value < 0.0001. **(B)** Root angles of the genotypes indicated 7h after the gravitropic stimulus. Numbers are individual seedlings. **(C)** Root bending kinetics of $P_{PIN2}$*:PIN1-2-1*, $P_{PIN2}$*:PIN2-1-2* or **(D)** $P_{PIN2}$*:PIN2-3-2* and $P_{PIN2}$*:PIN3-2-3* in *pin2* compared to wildtype and mutant. Insets show roots of the genotypes indicated 7 h after the gravitropic stimulus. Data points are mean and SE of n = 3 independent replicates consisting of 18-20 roots per genotype. Groups were compared by one-way ANOVA, followed by Dunnett post hoc test with Col-0 as control group. Significant differences are indicated by "*". n.s. not significant p-value ≥ 0.05, * p-value 0.01 to 0.05, ** p-value 0.001 to 0.01, *** p-value 0.0001 to 0.001, **** p-value < 0.0001.

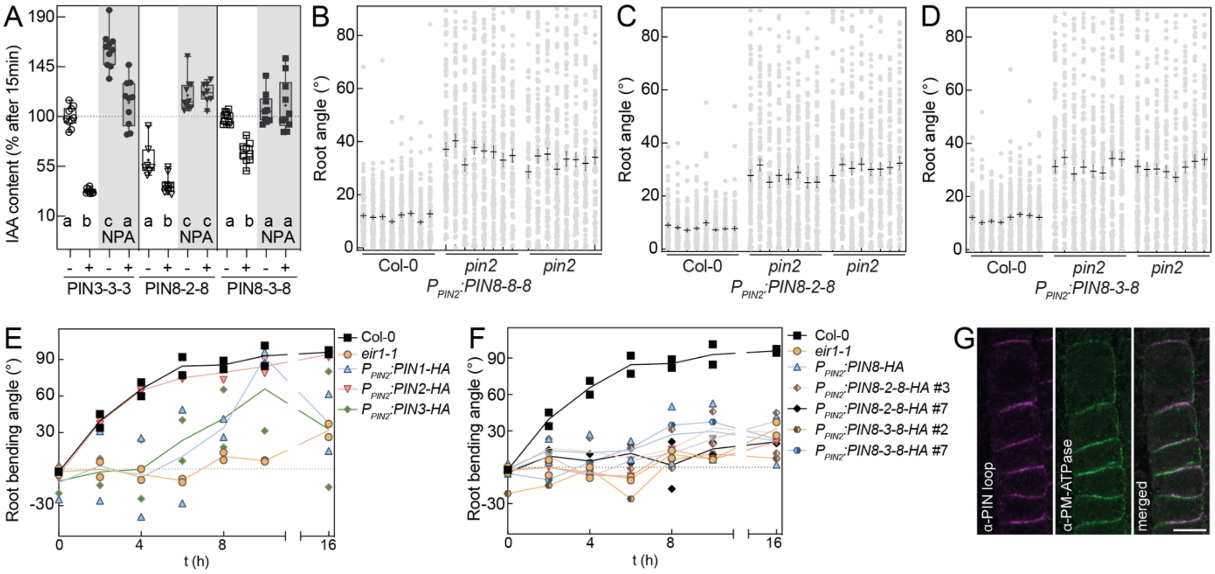


**Supplemental Figure 7. Properties of the PIN8 chimeras. (A)** PIN8 chimera are NPA-sensitive. PIN3 (circles), PIN8-2-8 (triangles) and PIN8-3-8 (squares) alone (-) or with PINOID (+) were expressed in oocytes. Oocytes were injected with radioactive IAA (empty symbols) or radioactive IAA + NPA (black symbols). The IAA content after 15 min of PIN3 was set to 100 % and the IAA content of the indicated samples at the endpoint of the experiment expressed relative to this. Data points represent individual oocytes (n = 7-10). Box plots ranges from 25th to 75th percentiles, whiskers mark the minimum and maximum values and the median is indicated. The mean is indicated by (+). Each PIN group (PIN3, PIN8-2-8, PIN8-3-8) was compared by one-way ANOVA followed by Tukey's posthoc test. Statistical groups are indicated by letters. **(B-D)** Identification of a segregating T2 line Data points are individual seedlings of transgenic lines (85 < n < 109) compared to wildtype (89 < 118) and mutant (78 < n < 108). The mean and SE are indicated. **(E)** Bending kinetics of two independent lines of *$P_{PIN2}$:PIN1-HA, $P_{PIN2}$:PIN2-HA* and *$P_{PIN2}$:PIN3-HA* compared to two lines of *pin2/eir1-1* and Col-0 Wildtype on 0.5 MS medium. n > 20 individual seedlings for each line. HA-tagged versions behaved comparable to the untagged versions shown in Fig. 4. The line indicates the mean of the two lines. **(F)** Bending kinetics of two independent lines of *$P_{PIN2}$:PIN8-2-8-HA* and *$P_{PIN2}$: PIN8-3-8-HA* compared to two lines of *pin2/eir1-1* and Col-0 Wildtype on 0.5 MS medium. n > 20 individual seedlings for each line. The line indicates the mean of the two lines. **(G)** Immunolocalization of PIN8-2-8-HA at the transition zone as specified by cell elongation PIN2 loop was decorated with a PIN2 rabbit antiserum (magenta), plasmamembrane ATPase is shown in green.
