## Supplementary material for "Transport properties of canonical PIN-FORMED proteins and the role of the loop domain in auxin transport": Janacek Extended Data Table 1

**Extended Data Table 1. Comparison of kinetic models to describe PIN-mediated IAA transport as a function of substrate concentration.** The data shown in Fig. 1A, D and G and Fig. 5B were used to compare the goodness of the fit of Michaelis-Menten model with a straight line. The preferred model is highlighted and Michaelis-Menten as preferred model and the corresponding $K_m$ value are shown in red.

| | PIN1 + D6PK | PIN1 + PID | PIN2 + D6PK | PIN2 + PID | PIN3 + D6PK | PIN3 + PID | PIN8 | PIN8-2-8 + PID |
|---|---|---|---|---|---|---|---|---|
| **Comparison of Fits** | Can't calculate | Can't calculate | Can't calculate | Can't calculate | Can't calculate | Can't calculate | Can't calculate | Can't calculate |
| **Null hypothesis** | Michaelis-Menten | Michaelis-Menten | Michaelis-Menten | Michaelis-Menten | Michaelis-Menten | Michaelis-Menten | Michaelis-Menten | Michaelis-Menten |
| **Alternative hypothesis** | Straight line | Straight line | Straight line | Straight line | Straight line | Straight line | Straight line | Straight line |
| **Conclusion (alpha = 0.05)** | Models have the same DF | Models have the same DF | Models have the same DF | Models have the same DF | Models have the same DF | Models have the same DF | Models have the same DF | Models have the same DF |
| **Preferred model** | Straight line | Straight line | Straight line | Straight line | Michaelis-Menten | Straight line | Michaelis-Menten | Straight line |
| **Michaelis-Menten** | | | | | | | | |
| **Best-fit values** | | | | | | | | |
| **Vmax** | 777.1 | Unstable | 106.5 | 423.9 | 2591 | Unstable | 616.8 | Unstable |
| **Km** | 238.3 | 2.503E+15 | 78.72 | 126.9 | 143.3 | 8.261E+14 | 48.85 | 9.786E+14 |
| **95% CI (profile likelihood)** | | | | | | | | |
| **Vmax** | 37.28 to +infinity | (Very wide) | ??? to +infinity | 29.17 to +infinity | 458.7 to +infinity | (Very wide) | 394.5 to 1542 | (Very wide) |
| **Km** | 4.983 to +infinity | ??? to +infinity | 0.7405 to +infinity | 3.059 to +infinity | 18.83 to +infinity | ??? to +infinity | 28.43 to 134.5 | -infinity to +infinity |
| **Goodness of Fit** | | | | | | | | |
| **Degrees of Freedom** | 16 | 16 | 16 | 16 | 16 | 16 | 13 | 14 |
| **R squared** | 0.7291 | 0.7644 | 0.3891 | 0.6244 | 0.9532 | 0.9874 | 0.9957 | 0.9408 |
| **Sum of Squares** | 758.6 | 1959 | 451.2 | 1199 | 3055 | 1195 | 78.21 | 4699 |
| **Sy.x** | 6.886 | 11.07 | 5.31 | 8.658 | 13.82 | 8.643 | 2.453 | 18.32 |
| **Constraints** | | | | | | | | |
| **Km** | Km > 0 | Km > 0 | Km > 0 | Km > 0 | Km > 0 | Km > 0 | Km > 0 | Km > 0 |
| **Straight line** | | | | | | | | |
| **Best-fit values** | | | | | | | | |
| **YIntercept** | 0.4874 | -0.3929 | 0.4369 | 0.6091 | 0.5294 | -2.334 | 1.264 | -1.328 |
| **Slope** | 3.076 | 5.425 | 1.156 | 3.04 | 16.97 | 20.83 | 10.59 | 21.62 |
| **95% CI (profile likelihood)** | | | | | | | | |
| **YIntercept** | -4.135 to 5.109 | -7.828 to 7.042 | -3.126 to 4.000 | -5.204 to 6.422 | -8.800 to 9.859 | -8.010 to 3.342 | -1.101 to 3.629 | -14.15 to 11.50 |
| **Slope** | 2.084 to 4.068 | 3.830 to 7.021 | 0.3918 to 1.921 | 1.792 to 4.287 | 14.97 to 18.98 | 19.61 to 22.05 | 10.03 to 11.14 | 18.52 to 24.73 |
| **Goodness of Fit** | | | | | | | | |
| **Degrees of Freedom** | 16 | 16 | 16 | 16 | 16 | 16 | 13 | 14 |
| **R squared** | 0.7299 | 0.7646 | 0.3911 | 0.6251 | 0.9528 | 0.988 | 0.9923 | 0.941 |
| **Sum of Squares** | 756.6 | 1958 | 449.7 | 1197 | 3083 | 1141 | 138.3 | 4682 |
| **Sy.x** | 6.877 | 11.06 | 5.302 | 8.649 | 13.88 | 8.445 | 3.262 | 18.29 |
| **Number of points** | | | | | | | | |
| **# of X values** | 24 | 24 | 24 | 24 | 24 | 24 | 32 | 32 |
| **# Y values analyzed** | 18 | 18 | 18 | 18 | 18 | 18 | 15 | 16 |
